# Comparative regulomics reveals pervasive selection on gene dosage following whole genome duplication

**DOI:** 10.1101/2020.07.20.212316

**Authors:** Gareth B. Gillard, Lars Grønvold, Line L. Røsæg, Matilde Mengkrog Holen, Øystein Monsen, Ben F. Koop, Eric B. Rondeau, Manu Kumar Gundappa, John Mendoza, Daniel J. Macqueen, Rori V. Rohlfs, Simen R. Sandve, Torgeir R. Hvidsten

## Abstract

Whole genome duplication (WGD) events have played a major role in eukaryotic genome evolution, but the consequence of these extreme events in adaptive genome evolution is still not well understood. To address this knowledge gap we used a comparative phylogenetic model and transcriptomic data from seven species to infer selection on gene expression in duplicated genes (ohnologs) following the salmonid WGD 80-100 million years ago. We find rare cases of tissue-specific expression evolution but pervasive expression evolution affecting many tissues, reflecting strong selection on maintenance of genome stability following genome doubling. Although ohnolog expression levels have evolved mostly asymmetrically, by diverting one ohnolog copy down a path towards pseudogenization, strong evolutionary constraints have frequently also favoured symmetric shifts in gene dosage of both copies, likely to achieve gene dose reduction while avoiding accumulation of ‘toxic mutations’. Mechanistically, ohnolog regulatory divergence is dictated by the number of bound transcription factors in promoters, with transposable elements being one source of novel binding sites driving tissue-specific gains in expression. Our results imply pervasive adaptive expression evolution following WGD to overcome the immediate challenges posed by genome doubling and to exploit the long-term genetic opportunities for novel phenotype evolution.

## Introduction

Whole genome duplication (WGD) events have played a major role in eukaryotic evolution by increasing genomic complexity and functional redundancy [1]. This can allow gene duplicates (referred to as ohnologs) to escape selective constraints and thereby accumulate previously forbidden mutations that may become adaptive [2]. In agreement with this idea, WGD has been associated with the evolution of adaptive traits in yeast [3], plants [4, 5], and vertebrates [6–8]. At the same time, it is also evident that most polyploids go extinct shortly after formation [9], and that becoming a successful new polyploid likely requires new adaptations to overcome fitness costs stemming from having a doubled genome [10, 11]. Yet, the importance of selection in shaping polyploid genome evolution in the aftermath of WGDs is still not well understood [1, 12].

Gene expression phenotypes are relatively easy to measure and compare, and represent a major source of complex trait variation [13] and novel adaptive phenotypes [14, 15]. Hence, there has been substantial interest in understanding consequences of WGDs on gene regulatory evolution. Comparative transcriptomics has both revealed immediate plastic responses to adjust gene dosages [16], as well as widespread regulatory divergence at evolutionary timescales [e.g. 17–20]. Ohnolog regulatory evolution is also mostly asymmetric, with one copy retaining an ancestral-like regulation, and the other copy losing and/or gaining expression in one or more tissue [12]. Although this observation can be reconciled with adaptive evolution of gene regulatory phenotypes following WGD, methodological limitations have made it difficult to distinguish between the outcomes of selection and neutral drift [12, 21].

Here we take a novel approach to improve our understanding of how selection shapes novel gene regulatory phenotypes following WGD. We first developed a flexible and user friendly version of a phylogenetic Ornstein-Uhlenbeck (OU) model of gene expression evolution [22, 23] in R (https://gitlab.com/sandve-lab/evemodel). The crux of this model is that it allows us to evaluate if changes in expression evolution deviate from the null hypothesis of stabilizing selection, and thereby identify putative adaptive shifts in expression regulation. We then used this model to analyze the liver transcriptome of four salmonids and three non-salmonid fish species to assess the impact of the 80-100 million year old salmonid fish specific WGD [24, 25]. We find that WGD leads to a burst of gene expression evolution, leading to rare tissue-specific gains in expression and pervasive tissue non-specific dosage selection, reflecting both adaptive possibilities afforded by genome doubling and immediate challenges that must be overcome to succeed as a polyploid lineage.

## Results

### Adaptive shifts in expression levels following WGD

To study expression level evolution following WGD we generated RNA-seq datasets from livers (four biological replicates) of four salmonids and three non-salmonid outgroup species (Figure 1A). We then computed gene trees to identify retained ohnologs from the salmonid WGD. In total, we included 10,154 gene trees in our analyses (Supplementary figure 1), of which sixty-five percent (6689 trees) contained ohnologs derived from the salmonid WGD. For each gene tree we then applied a phylogenetic Ornstein-Uhlenbeck (OU) process model to test for adaptive shifts in expression evolution (referred to simply as ‘shifts’) in the ancestor of the salmonids included in this study (Figure 1B, Supplementary figure 2, 3 and 4).

**Figure 1.**
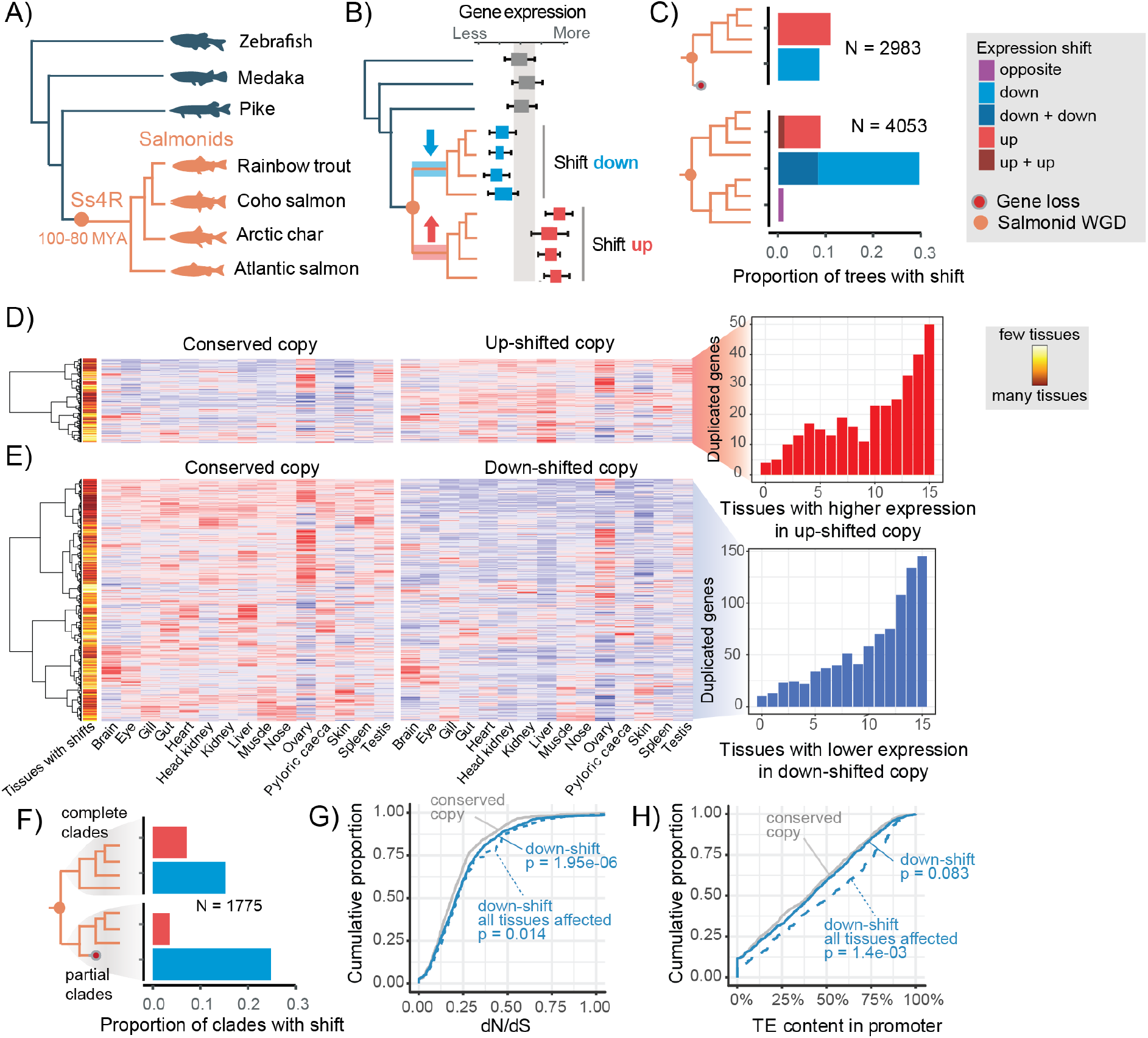
Expression level evolution following WGD. (A) Phylogenetic tree of the species included in the study, with the estimated time of the salmonid-specific whole genome duplication (Ss4R) indicated. (B) Conceptual illustration of the expression level evolution tests. (C) Proportion of complete singleton (top) and ohnolog (bottom) gene trees with significant shifts in expression level in a salmonid ancestor. (D and E) Heatmaps show tissue expression, from an independent tissue atlas in Atlantic salmon, of ohnolog pairs where one copy has shifted up (D) or down (E) in liver. Barplots show the distribution of the number of tissues where the shifted copy has lower or higher expression than the conserved copy. Only ohnologs from complete orthogroups (panel C) are included in the heatmap. Each ohnolog pair (row) is scaled so that red signifies the highest expression across the two copies and blue the lowest. The color bar indicates the number of tissues that are experiencing a shift in expression in the same direction as that of liver (down (D), up (E)) between the shifted and conserved copy. (F) Proportion of partial gene trees (i.e. trees with some gene loss) with significant shifts in expression level in a salmonid ancestor. The shadings indicate that we report here up/down shifts for the complete salmonid clade and the partial salmonid clade separately, which is in contrast to panel C where both salmonid clades are complete and therefore indistinguishable. (G) Cumulative proportion of dN/dS for ohnologs with one copy shifted down, versus their conserved counterpart. Results are shown for all ohnologs with one copy shifted down (down-shift) and for the subset that is down-shifted in all tissues in the tissue atlas (down-shift all tissues affected). (H) Cumulative proportion of TE content in promoters of ohnologs with one copy shifted down.

**Figure 2.**
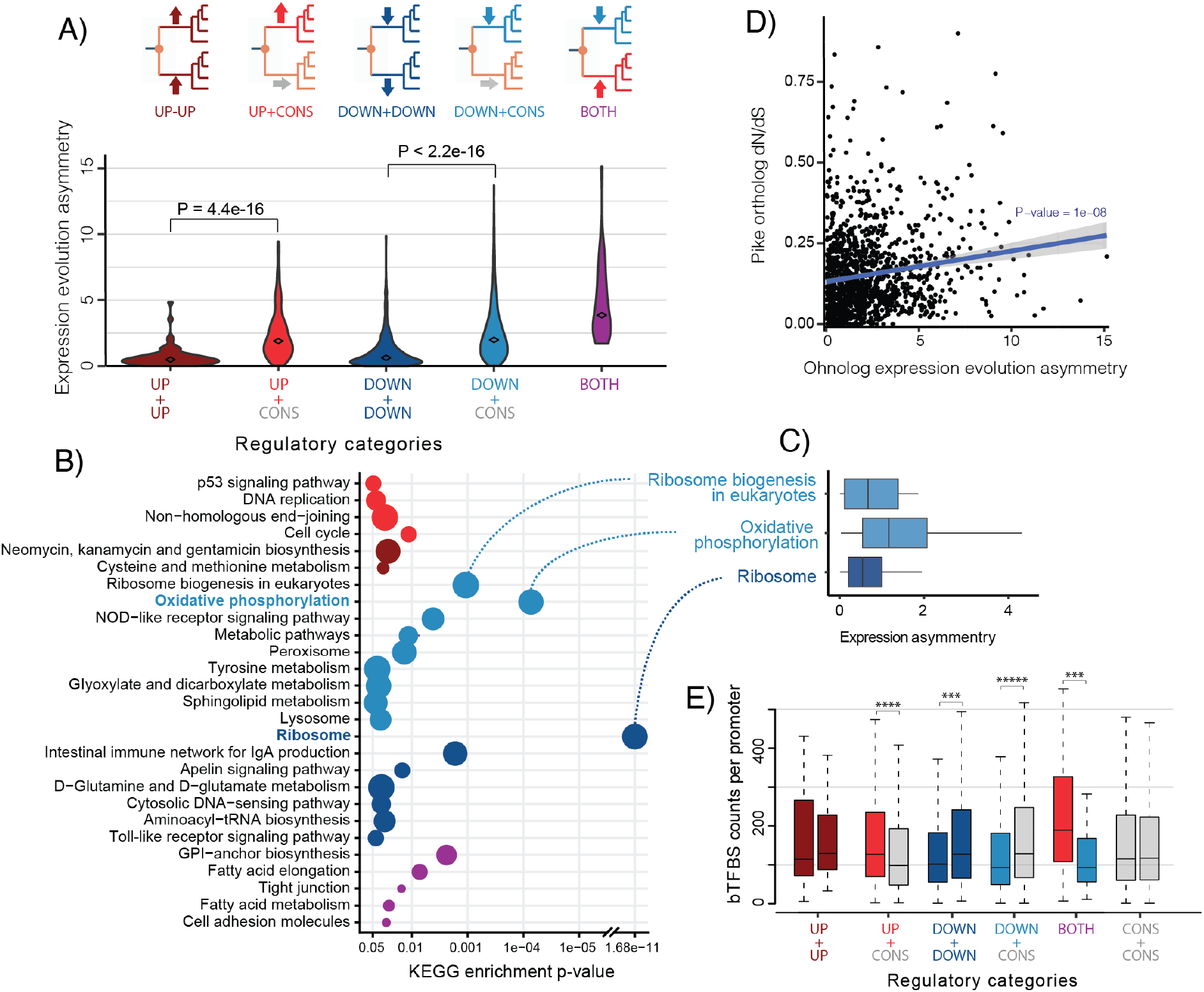
Symmetry of regulatory divergence. (A) Ohnolog expression evolution categories and expression evolution asymmetry for ohnologs in each evolutionary category. The expression asymmetry is calculated as the absolute value of the mean difference between ohnolog pair expression levels in all salmonid species. One sided Wilcoxon test p-values are reported for significant asymmetry differences between symmetric and asymmetric regulatory categories. (B) KEGG pathways significantly enriched (p < 0.05) in different expression evolution categories. Larger circles indicate a higher proportion of genes in the pathway with the shift. (C) Expression asymmetry between salmonid ohnolog pairs in selected pathways, calculated by taking the absolute value of the mean difference in expression between ohnolog pairs in all salmonid samples. (D) Correlation between expression asymmetry (see (C) for details) and the dN/dS of the ortholog in the pike sister lineage. (E) Predicted bound TFBS from TF-footprinting in promoters of ohnologs in the five expression evolution categories as well as those ohnologs with no significant shift in expression levels. For each ohnolog pair in each category, copies are grouped based on the lowest (to the left) and highest (to the right) p-value in the OU-test for expression level shift. P-values from significant paired Wilcoxon tests are indicated above boxplots: *** < 1e-03, **** < 1e-04, ***** = 0.

**Figure 3.**
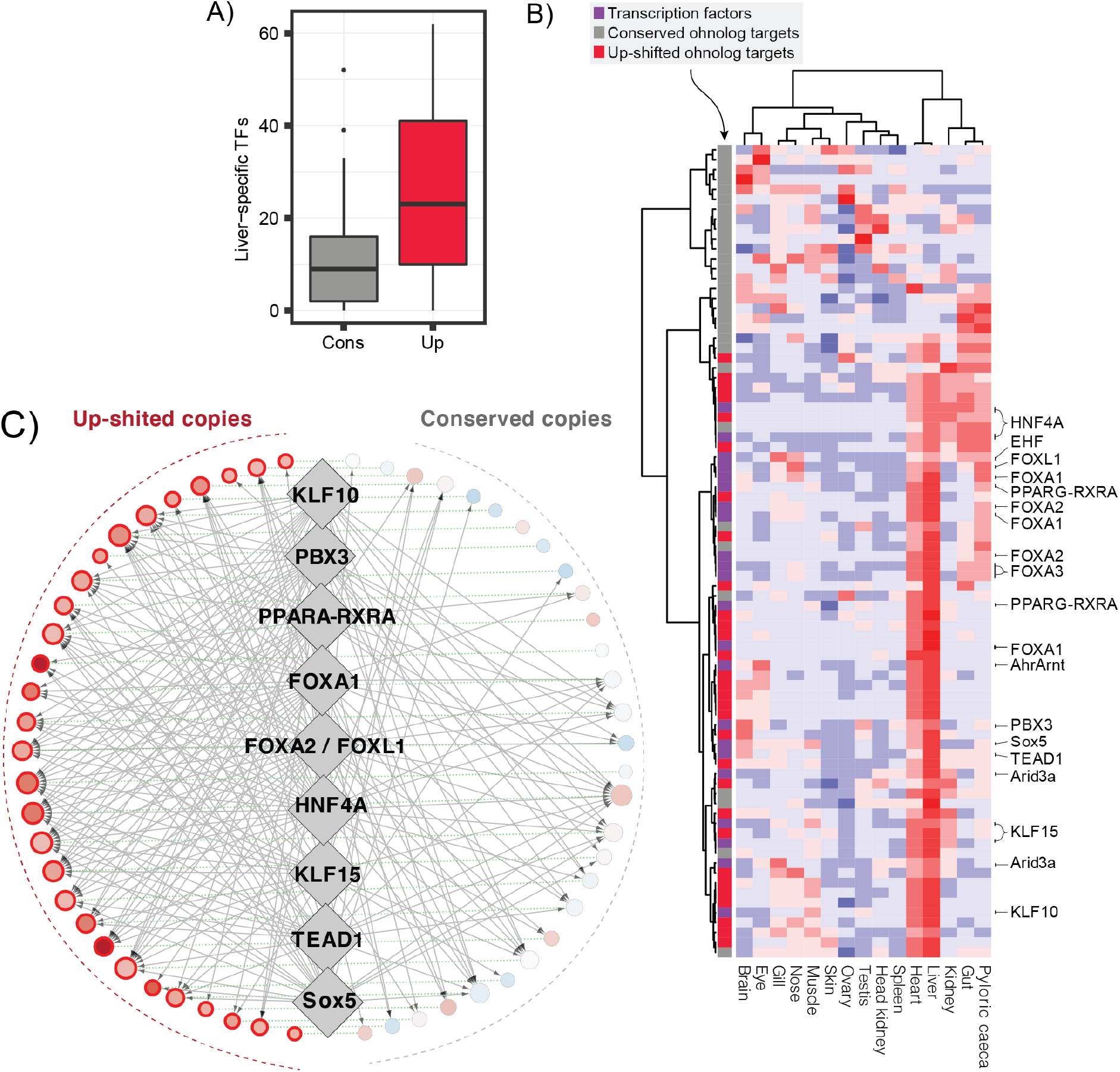
Transcription factor binding site evolution. (A) The number of liver-specific TFs (56 in total) with at least one bTFBS in the promoters of the 30 ohnologs with one liver-specific up-shifted copy (Up) or one conserved copy (Cons). (B) Tissue expression of the 30 ohnolog pairs where one copy has evolved a liver-specific gain in expression (color bar: up-shifted copies are red and conserved copies are grey) and 22 liver-specific TFs predicted to bind at least one-third of the targets (purple). TFs are named according to their motif(s) in JASPAR. Liver-specific genes are defined as having liver expression levels in the 90% quantile and tau-scores > 0.6. Each gene (row) is scaled so that red signifies the highest expression across the tissues and blue the lowest. (C) Regulatory network reconstructed for the ohnologs and selected TFs from B) using footprinting data. Ohnologs are represented by circles sized by their regulatory complexity (in-degree) and colored according to their evolutionary expression shift with red signifying up-shift and blue down-shift. TFs are represented by diamonds with the nine most up-shift-biased TFs shown. A directed grey edge means that the TF has at least one bTFBS in the promoter of the gene. A dotted undirected green edge connects ohnologs.

Two major observations arise from this analysis. First, it is evident that the rate of adaptive gene expression evolution is increased for salmonid ohnologs. Forty percent of trees (1649) with retained ohnologs display evolution of novel expression levels in at least one ohnolog compared to only twenty percent of trees with a single copy gene (Figure 1C). Secondly, there is a clear difference in the nature of the expression evolution between ohnologs and singleton genes. Ohnologs are strongly biased towards evolving decreased expression levels following WGD (Figure 1C), with 75% (1234/1649) of the ohnolog pairs displaying a shift down in either one or both copies. Conversely, singletons show a small bias towards evolving increased expression (Figure 1C). This difference could not be explained by differences in statistical power related to systematic differences in gene expression levels between singletons and ohnologs (Supplementary figure 5).

To test if the identified expression level shifts following WGD were tissue-specific, we analyzed RNA-seq data from 15 Atlantic salmon tissues (Supplementary figure 6A). We find that most cases of expression evolution are not liver-specific (Figure 1D-E), and that this is true both for genes evolving increased and decreased expression following WGD. When one ohnolog copy had evolved a shift in liver expression level, this copy also displayed similar trends in the majority of the other 14 tissues compared to it’s conserved ohnolog partner (shift down: 77% (682/885), shift up: 70% (221/317)). Hence, evolution of liver-specific changes in ohnolog expression following WGD is rare, irrespective of the directionality of change.

Upon reaching a new optimal ohnolog gene dosage, the expectation is that the copy with highest expression level (i.e. conserved copy) contributes the most to the proteome and cell function, which will result in reduced purifying selection pressure on the evolved copy (i.e. down-shifted copy) [26]. Several lines of evidence support this expectation. Firstly, species-specific gene loss events (expected for genes evolving under relaxed selection) are associated with increased probability of evolving lower liver expression in one copy (Figure 1F) and with increased probability of the down-shifted copy to have reduced expression levels across all the other 14 tissues (Fisher’s exact test, p = 3.1e-07, Supplementary figure 6B). Secondly, we find that the down-shifted copy shows increased signatures of relaxed purifying selection on coding sequences in the form of elevated dN/dS rates (Figure 1G, p = 2.1e-6, N = 732, one-sided paired Wilcoxon test, Supplementary figure 7). Lastly, we also observe that down-shifted ohnolog copies have a significantly higher load of potentially destructive transposable element (TE) insertions in promoters compared to the conserved partner (Figure 1H, one-sided paired Wilcoxon test, p = 6.5e-4, Supplementary figure 8). Importantly, the effect size of increased dN/dS and TE-load were larger when only considering ohnologs with signatures of down-shift across all tissues (Figure 1G-H).

Pervasive differences in purifying selection pressure within individual ohnolog pairs raise the question of whether these ohnologs might belong to duplicated genome blocks experiencing large-scale biases in selective constraints (known as biased fractionation). In line with previous studies on teleosts (Conant 2019; Xu et al. 2019) we found significant biases in gene loss, albeit only in 9 of 47 syntenic duplicate blocks. However, we did not find equivalent large-scale biases in expression loss (Supplementary figure 9), thus rendering regional differences in selection constraints an unlikely explanation for the large number of ohnologs experiencing loss of expression in one copy.

In conclusion, we find widespread signatures of adaptive regulatory evolution in retained ohnologs following WGD, however most adaptive events were associated with ohnolog gene dose reduction across many tissues. Thus, ohnolog copies that evolve lower expression levels compared to their partner continue to evolve under relaxed purifying selection pressure, following a likely path towards pseudogenization.

### Strong selection on housekeeping gene dose after WGD

To test if selection on gene regulation following WGD was linked to particular cellular functions or pathways we performed KEGG enrichment analyses for two ohnolog gene sets that had evolved either increased (up) or decreased (down) expression levels. Genes with increased expression level were enriched (Fisher’s exact test, p < 0.05) in three pathways; ‘fatty acid elongation’, ‘fatty acid metabolism’, and the ‘cell cycle’ (Supplementary table 1). Detailed analysis identified 29 up-shifted genes encoding proteins with essential cell division functions. These genes were highly enriched in protein-protein interactions conserved in both unicellular and multicellular eukaryotes (Supplementary table 2, Supplementary figure 10), which suggests compensatory adaptation to maintain genomic integrity by increased gene dosage.

Down-shifted genes had comparatively stronger functional signatures (Supplementary table 1) with nine enriched pathways (Fisher’s exact test, p < 0.05). The three pathways with the strongest enrichment were ‘oxidative phosphorylation’ (p = 0.003) involved in mitochondrial-associated cellular energy production, ‘ribosome biogenesis in eukaryotes’ (p = 0.008) which consists of genes involved in assembly of the ribosome, and ‘ribosome’ (p = 5.6e-9) which consists of ribosomal subunit genes (Supplementary figures 11, 12 and 13). These results support strong selection on gene dosage for many housekeeping functions following WGD, which aligns well with our observation (Figure 1D-E) that most expression level shifts occurred across most tissues.

The gene balance hypothesis predicts that selection operates to maintain stoichiometry of interacting gene products [27], and this is believed to result in long term retention of ohnologs. Using the human orthologs of salmonid genes we queried the CORUM database of protein complexes and found that the proportion of ohnologs in protein complexes was only slightly higher (28%) than the proportion of singletons (22%) (Fisher’s exact test, p = 1.04e-5, Supplementary figure 14). It is also plausible that stoichiometric imbalances could be rescued through evolution of novel gene dosage. Under this model we predict that singletons in protein complexes that contain ohnologs should be enriched for shifts up in expression, while shifts down are predicted for ohnologs in complexes with singletons. These predictions are not well supported for singletons (Fisher’s exact test, p = 0.07) nor ohnologs (Fisher’s exact test, p > 0.48) (Supplementary table 3).

Taken together, although we find strong evidence for dosage selection in general, we do not find support for selection on protein-complex stoichiometry (relative dosage) playing a major role in ohnolog retention or regulatory evolution following WGD.

### Mechanism driving ohnolog regulatory divergence is associated with functional constraints

Our analysis allows us to assign ohnolog pairs to different categories (Figure 2A) that potentially represent distinct evolutionary routes to new gene dosage optimums after WGD. Indeed our results show that ohnolog pairs with expression evolution shifts in the same direction evolve more symmetrically (down+down and up+up) while ohnologs where expression shifts occur in only one copy or in opposite directions display stronger asymmetric divergence (e.g. up/down+conserved) (Figure 2A). To explore the links between these modes of regulatory divergence and gene function we performed KEGG enrichment on each expression evolution category. Twenty-seven pathways were found enriched across these categories (Figure 2B, Supplementary table 4), which is more than twice as many as when grouping ohnologs into up- or down-shifted genes (Supplementary table 1). This supports that different pathways are biased towards either symmetric or asymmetric regulatory evolution. The three most enriched pathways were the same as when testing up- and down-shifted genes only, but our stratification on regulatory categories of ohnologs reveals that ribosomal subunit ohnologs (‘Ribosome’) evolved lower gene dosage through highly symmetrical down-shifts, while ‘oxidative phosphorylation’ and ‘ribosome biogenesis in eukaryotes’ are biased towards asymmetric divergence (Figure 2C). As ribosome subunit genes are known to be extremely slowly evolving genes (i.e. high sequence evolution constraints) we tested whether there is a broader correlation between sequence constraints and regulatory symmetry. Indeed, we find that ohnologs expression level symmetry is significantly correlated with the level of purifying selection on coding sequences (Spearman correlation, p = 1e-8, Figure 2D).

To further dissect regulatory mechanisms driving ohnolog expression level evolution, we generated high coverage ATAC-seq data from the liver of Atlantic salmon and identified bound transcription factor binding sites (bTFBSs) using a footprinting approach (Supplementary figure 15). We hypothesized that ohnolog regulatory evolution symmetry is shaped by the relative importance of selection on cis-versus trans-mutations. One simple prediction from this is that ohnolog pairs where one copy has evolved novel expression would have higher promoter divergence than ohnolog pairs with symmetric evolution. The divergence of bTFBSs in promoters (−3000/+200bps from transcription start site) largely matched this prediction (Figure 2E) with ohnologs having more asymmetric expression shifts (up+cons and down+cons) differing more with respect to the number of bTFBSs in their promoters compared to symmetrically evolving ohnologs (up+up, down+down and cons+cons) (Figure 2E). This offers a simple explanation of expression divergence after WGD, where genes with decreased expression level have lost TFBSs, and genes with increased expression have gained TFBSs, compared to the ancestral promoter structure. Comparing the overall similarity of promoters, computed as the correlation of bTFBS between symmetrically evolving (down+down) and asymmetrically evolving (down+cons) ohnolog pairs, did not reveal a similar trend (Wilcoxon test, p = 0.234, Supplementary figure 16), which is consistent with high turnover of bTFBS even for highly conserved genes [28].

Together these results support that evolutionary constraints at the coding sequence divert ohnologs down different evolutionary routes towards novel gene dosage - either in an asymmetric or symmetric fashion.

### Adaptive gain in liver expression through aquisition of tissue-specific cis-regulatory elements

Although the vast majority of adaptive expression evolution was associated with selection on lower gene dosage, our OU-analyses did reveal 30 ohnolog pairs where one copy had evolved liver-specific adaptive gains in expression following WGD. These genes are predicted to be involved in a variety of functions such as developmental processes, cell fate specificity, as well as more liver-centric functions such as endocrine signalling, lipid- and fatty-acid metabolism (Supplementary table 5). To better understand the regulatory mechanisms involved in the evolution of these potential novel liver functions, we used our TF-footprinting data to test the hypothesis that adaptive gains in liver expression are linked to the acquisition of binding sites for TFs controlling liver-specific regulatory networks. Indeed, we found that promoters of up-shifted copies were occupied by many more liver-specific TFs than their non-shifted partners (Figure 3A, Wilcoxon paired test, p = 7.7e-05). These liver-specific TFs are thus candidates for being involved in regulatory rewiring of up-shifted ohnologs (Figure 3B). Interestingly, many TFs with the strongest bias towards occupying the promoters of up-shifted ohnolog copies have known general liver functions (i.e. hepatocyte nuclear factors; FOX1A, HNF4A) [29] and roles in lipid metabolism (RXR, PPARG, KLF15) [30, 31] (Figure 3C, see Methods for details).

Next, we hypothesized that liver-specific increases in expression are driven by gains in new TFBSs. One way promoters can gain novel TFBSs is through insertions of TEs that either contain a functional TFBS or subsequently accumulate mutations that give rise to new TFBSs [32]. Indeed, we did find that TFBSs predicted to be bound by liver-specific TFs overlapped TEs more often in up-shifted copies than in conserved copies (Wilcoxon paired test, p = 0.037, Supplementary figure 17A). Furthermore, at the level of TE superfamilies we found that the TIR TC1-Mariner TE superfamily were associated with gain in liver-specific bTFBS in up-shifted copies (p = 0.018, Supplementary figure 17B), which included known liver and lipid metabolism transcription factors such as HNF4A, KLF15 and RXRA (Supplementary table 6).

In conclusion, we find that adaptive gain in liver-specific expression is strongly associated with gain in liver-specific bound TFBSs, some of which have been facilitated by transposable element insertions.

## Discussion

The consequence of WGDs for evolution of novel adaptations, including gene expression phenotypes, has been an actively debated topic within evolutionary biology [1]. A key challenge has been to distinguish neutral from adaptive evolution in systems where experimental evolution is not possible [12]. Here, we generated a large comparative transcriptomics dataset, and for the first time applied a formal phylogenetic model to infer selection on gene expression in the aftermath of a vertebrate WGD that occured 80-100 million years ago.

### Selection on gene dosage ameliorates immediate polyploid fitness costs

Newly formed polyploids often display augmented rates of abnormal mitosis, chromosome loss and gross chromosomal rearrangements (Storchová and Pellman, 2004; Storchová et al. 2006). Hence, a primary challenge for the evolutionary success of polyploids is to maintain genomic stability. In line with this, we find that adaptive evolution of gene expression was highly biased towards cellular functions not specific to the liver (Figure 1E-F, Figure 3B) and with a clear potential impact on genome stability. Firstly, we find genes directly involved in the cell cycle to be enriched for adaptive evolution (higher dosage). Related genes have experienced selective sweeps following WGD in plants [33, 34]. Furthermore, we find strong evidence for selection on genes involved in oxidative phosphorylation (lower dosage). Polyploidization in plants, fungi, and mammalian cells have been shown to increase levels of reactive oxygen species, which is causally linked to increased cellular stress, cell cycle failure, and increased genome instability [35–37]. Lastly, we find adaptive expression evolution (lower dosage) for genes involved in translation (ribosome subunits and ribosome assembly) after WGD. Regulation of translation also interacts with cell cycle regulation, with potential implications for genome stability [38]. However, selection for decreased expression of translation-related genes could also be linked to direct fitness costs of wasteful protein translation or harmful effects linked to the over-production of particular proteins. Overall, our study provides evidence for a scenario where a critical first step in becoming a successful polyploid lineage is pervasive adaptive evolution on gene dosage to ameliorate fitness costs linked to genome stability.

### Long term ohnolog retention and selection on gene dosage

Following initial selection on gene dosage, long term retention of ohnologs could be driven by various adaptive processes [21, 39], including adaptive regulatory evolution. We find that positive selection on novel tissue-specific regulatory functions (Figure 1C: up+cons) is rare and likely contribute little to the total number of retained ohnologs. Furthermore we observe evidence for selection on stoichiometry (Supplementary figure 14), but little support for adaptive expression evolution to rescue stoichiometric imbalances (Supplementary table 3). Finally, we find significant correlation between constraints at the coding sequence level and symmetry of regulatory evolution (Figure 2D). One potential explanation for this pattern could be the ‘toxic effects model’ where long term conservation of ohnologs is intrinsically linked to the ‘danger’ of accumulating highly toxic coding sequence mutations [40, 41]. We therefore hypothesize that in situations where lowering total gene dosage increases fitness, but the tolerance for accumulation of deleterious mutations is low (i.e. the toxic effect), symmetric ohnolog evolution towards lower gene dosage could be favoured over slow pseudogenization of one copy.

Regulatory divergence after gene duplication is hypothesized to be linked to evolution of local chromatin landscapes [18, 42]. Using ATAC-seq data we show that signals of adaptive expression level shifts are associated with the numbers of bound TFBSs (Figure 2E), consistent with a billboard-like model of gene regulation [43]. Furthermore, we find that both loss of expression (Figure 1H) and tissue specific gains in expression level (Supplementary figure 17) is linked to TE activity, highlighting the dual role of TEs in regulatory evolution following WGD.

### Conclusion

Our study supports pervasive selection on gene dosage across millions of years following WGD, in particular for genes involved in basic cellular maintenance and genome stability. Interestingly, many of the same genes and pathways also show similar responses in gene dosage adjustments immediately after polyploidization in plants [16]. Reconciling these immediate effects of polyploidization with our findings strongly supports the following model: Plastic genome regulatory response to polyploidization alleviate immediate fitness costs following genome doubling. Since gene loss is absent in early generations polyploids, the initial selection pressure on gene regulatory phenotypes is likely a result of selection on absolute dosage, rather than stoichiometry. Over evolutionary time-scales however, selection will favour and fix regulatory mutations that can ‘hard code’ novel transcriptional phenotypes to optimize gene dosages (as seen following the salmonid WGD). Together, this paper points to critical genome regulatory adjustments for becoming a successful polyploid lineage.

## Methods

### Ortholog inference

For ortholog inference we used thirteen species including six salmonids (*Thymallus thymallus, Hucho hucho, Salmo salar, Salvelinus alpinus, Oncorhynchus mykiss* and *Oncorhynchus kisutch*), four telosts as outgroups to the salmonids (*Danio rerio, Oryzias latipes, Gasterosteus aculeatus* and *Esox lucius*), one non-teleost fish (*Lepisosteus oculatus*) and two mammals as outgroups to the teleosts (*Homo sapiens* and *Mus musculus*). Protein sequences were obtained from ENSEMBL (release 92) for *H. sapiens, M. musculus, L. oculatus, D. rerio, O. latipes* and *G. aculeatus*, from NCBI RefSeq assemblies for *S. salar* (GCF_000233375.1), *S. alpinus* (GCF_002910315.2), *O. mykiss* (GCF_002163495.1), *O. kisutch* (GCF_002021735.1) and *E. lucius* (GCF_000721915.3), from the genome paper for *T. thymallus* [44] and from an in-house annotation using Transdecoder (https://github.com/TransDecoder/TransDecoder/wiki) for *H. hucho* (GCA_003317085). The single longest protein per gene was assigned to gene ortholog groups (orthogroups) using OrthoFinder (v2.3.1) [45]. For each orthogroup, the corresponding CDS sequences were aligned using MACSE (v2.03) before gene trees were generated and reconciled against the species tree using TreeBest (v1.9.2). The gene trees were then split at the level of monophyletic teleost clades, defining what we refer to as trees in this article, and again at the level of the salmonid clade (excluding *T. thymallus* and *H. hucho*), defining the Ss4R duplicate clades. Trees were then selected based on their topology (Supplementary figure 1). Specifically, this filtered any trees that showed more than two salmonid clades or that contained additional paralogs inside the salmonid clades or in the outgroup species. Trees with all orthologs retained in the salmonid clade(s) were designated as complete, and otherwise as partial. In addition, trees were excluded from further analysis if (1) one or both salmonid clades had no expressed genes (zero mapped reads, RNA-seq data described below), (2) the *E. lucius* ortholog was missing or not expressed and (3) both the *D. rerio* and *O. latipes* orthologs were missing or not expressed.

### RNA-sequencing data

Liver tissue samples were collected from adult individuals of *D. rerio* (zebrafish), *O. latipes* (medaka), *E. lucius* (pike), *O. mykiss* (rainbow trout), *S. alpinus* (Arctic char), and *O. kisutch* (coho salmon) (Figure 1A). Samples were taken in replicates of four, or three in the case of rainbow trout. All fish were raised in fresh water under standard rearing conditions in aquaculture facilities (salmonids), animal laboratory facilities (zebrafish and medaka), or restocking hatcheries (pike). Total RNA was extracted from the liver samples using the RNeasy Plus Universal Kit (QIAGEN). Quality was determined on a 2100 Bioanalyzer using the RNA 6000 Nano Kit (Agilent). Concentration was determined using a Nanodrop 8000 spectrophotometer (Thermo Scientific). cDNA libraries were prepared using the TruSeq Stranded mRNA HT Sample Prep Kit (Illumina). Library mean length was determined by running on a 2100 Bioanalyzer using the DNA 1000 Kit (Agilent) and library concentration was determined with the Qbit BR Kit (Thermo Scientific). Paired-end sequencing of sample libraries was completed on an Illumina HiSeq 2500 with 125-bp reads. Raw RNA-seq and processed count data have been deposited into ArrayExpress under the projects E-MTAB-8959 and E-MTAB-8962. For *S. salar* (Atlantic salmon), RNA-seq data was obtained from a feeding trial using four samples from individuals in freshwater fed a marine based diet [46], available in the European Nucleotide Archive (ENA) under project PRJEB24480 (samples: ERS2101563, ERS2101567, ERS2101568, ERS2101569).

To generate gene expression data, RNA-seq reads were mapped to the annotated reference genomes using the STAR aligner with default settings [47]. RSEM [48] was used to estimate read counts and Transcripts Per Million reads (TPM)-expression values that are normalized for average transcript lengths and the total number of reads from each sample.

The trimmed mean of M values (TMM), from the R package edgeR [49], was used to compute normalization factors for the gene expression data. The replicates were first normalized within each species and then between species (Supplementary figure 2). Between-species normalization was accomplished by first computing species-specific normalization factors using genes from singleton orthogroups (i.e. groups containing only one gene from each species) and their mean expression values (i.e. mean of the replicates within each species), and then by normalizing the individual replicates from each species using these normalization factors. All expression values were log transformed (log2(TPM+0.01)) prior to testing for expression shifts.

### Evolutionary shifts in gene expression

The EVE model [22] was used to test for shifts in gene expression levels in the salmonid clade(s) within each gene tree. For this paper, we developed and implemented a user friendly version of the EVE algorithm in R (https://gitlab.com/sandve-lab/evemodel). This method models an OU process, i.e. random drift in expression level that is constrained around an optimal level. The test compares a model with two optimal expression levels, one for the salmonid branch and another for the outgroup species, against the null-model which has the same optimal expression level across the entire tree (Supplementary figure 3C). For ohnolog gene trees which contain two duplicate salmonid clades, each clade was tested separately by removing the other salmonid clade.

EVE was given the expression data for each species (four samples/replicates per species) and the species tree produced by OrthoFinder. For every ortholog, a likelihood ratio test (LRT) score is calculated, representing the likelihood of the alternative hypothesis over the null hypothesis. LRT scores were compared to a Chi squared distribution with one degree of freedom and scores above the 95% quantile were considered to be significant. EVE reports estimates of the expression optimum for the salmonid branch and the rest of the tree (i.e. outgroup species), and the difference between salmonid estimates and outgroup estimates provided the direction of the expression shift.

### Tissue atlas

Gene expression data from an Atlantic salmon tissue atlas [17] was clustered using Pearson correlation and the R function hclust with method = “ward.D”. Heatmaps were drawn using the R function pheatmap with scale = “row”.

### Coding sequence selection pressure

We estimated branch specific selection pressure on coding sequences in ohnolog gene trees by calculating dN/dS measured at the branch from the WGD node to the root of each duplicate clade using the aBSREL (adaptive Branch-Site Random Effects Likelihood) method [50] in Hyphy (Hypothesis Testing using Phylogenies) [51]. A one-sided paired Wilcoxon test was then performed to test if there is a difference in selection pressure between ohnolog pairs classified as asymmetrically shifted at the expression level.

### Transposable elements

Transposable element (TE) annotations were taken from [17]. For Atlantic salmon genes, we calculated the proportion of gene promoter sequence (+2kb/-200b from TSS) that was overlapped with TEs using bedtools intersect of promoter and TE annotations. We used a one-sided paired Wilcoxon test to test the hypothesis that, for ohnologs with an asymmetric shift down in expression, the shifted copy had a higher proportion of TE overlap than the conserved copy.

### Gene function enrichment

We assigned KEGG pathway annotations to the orthogroups based on the Northern pike ortholog and it’s KEGG annotations. We then tested each set of ohnologs within an expression shift category for the enrichment of KEGG pathways using the kegga function from the R package limma, with all tested ohnologs as the background.

### Protein complexes

We assigned orthogroups as being in a protein complex or not based on the human ortholog and it’s protein complex annotations from the CORUM database [52]. We used the Fisher’s exact test, for singleton and ohnolog genes, to test whether more genes within an expression shift category were in a protein complex than expected by chance.

### ATAC-seq generation and TF footprinting

Four Atlantic salmon (freshwater stage, 26-28g) were euthanized using a Schedule 1 method following the Animals (Scientific Procedures) Act 1986. Around 50mg homogenized brain and liver tissue was processed to extract nuclei using the Omni-ATAC protocol for frozen tissues [53]. Nuclei were counted on an automated cell counter (TC20 BioRad, range 4-6 um) and further confirmed intact under microscope. 50,000 nuclei were used in the transposition reaction including 2.5 µL Tn5 enzyme (Illumina Nextera DNA Flex Library Prep Kit), incubated for 30 minutes at 37 °C in a shaker at 200 rpm. The samples were purified with the MinElute PCR purification kit (Qiagen) and eluted in 12μL elution buffer. qPCR was used to determine the optimal number of PCR cycles for library preparation [54] (8-10 cycles used). Sequencing libraries were prepared with short fragments and fragments >1,000 bp removed using AMPure XP beads (Beckman Coulter, Inc.). Fragment length distributions and confirmation of nucleosome banding patterns were determined on a 2100 Bioanalyzer (Agilent) and the library concentration estimated using a Qubit system (Thermo Scientific). Libraries were sent to the Norwegian Sequencing Centre, where paired-end 2 x 75 bp sequencing was done on an Illumina HiSeq 4000. The raw sequencing data for brain and liver is available through ArrayExpress (Accession: E-MTAB-9001).

Reads were mapped using BWA-MEM [55]. Duplicate reads and reads mapping to mitochondrial or unplaced scaffolds were removed. Peaks were called using MACS2 [56]. TF footprinting was performed with TOBIAS [57] based on the aligned reads, peaks and TF motifs from JASPAR (JASPAR 2020 non-redundant vertebrate CORE PFMs) [58]. TOBIAS performs Tn5 bias correction, generates footprint scores for each base within the peaks, scans for TFBSs using the given TF motifs, and finally classifies each TFBS as bound or unbound based on the footprint scores.

For the analysis of ohnolog pairs with evolved liver-specific expression increases in one copy, we identified 30 up+cons pairs (60 target genes) where the liver expression of the up-copy was at least 90% of the maximum expression in the tissue atlas and the up-copy had a tissue specificity score (tau) > 0.6 [17]. To identify regulators of these genes, we BLASTed UniProt TF sequences with a motif in JASPAR to the Atlantic salmon proteome, and retained the top four hits with E-value < 1E-10 and alignment length > 100. We then filtered these TFs for having bTFBS in the promoter of at least 20 of the target genes and for having liver-specific expression (same criteria as for up-targets). This resulted in 22 liver-specific TFs predicted to bind 17 different JASPAR motifs in 52 target promoters (Figure 3B-C). Finally, to draw the network in Figure 3C we (1) selected, for each JASPAR motif, the single TF with the strongest evolutionary shift in expression, (2) removed JASPAR motifs with highly similar binding profiles (>80% overlap in target genes, retaining the TF with the strongest evolutionary shift), (3) merged TFs associated with more than one JASPAR motif into one node and selected the nine TFs with the strongest bias towards up-shifted targets.

### Reproducibility

The scripts developed to implement analyses described in this study are available here: https://gitlab.com/sandve-lab/gillard-groenvold

## Declarations

### Ethics approval and consent to participate

The fish used in this study were treated according to the Norwegian Animal Research Authority (NARA) in accordance with the Norwegian Animal Welfare Act of 19th of June 2009.

### Availability of data and materials

Raw RNA-seq and processed count data have been deposited into ArrayExpress under the projects E-MTAB-8959 and E-MTAB-8962. Raw ATAC-seq data is also available through ArrayExpress under project E-MTAB-9001. The scripts developed to implement analyses described in this study are available at https://gitlab.com/sandve-lab/gillard-groenvold.

### Authors’ contributions

SRS, TRH and RR conceived the study. GG, LG, LR, MMH, ∅M, MKG, DJM, JM, BK, and EBR were involved in formal generation, analysis and/or curation of data. LG, GG, SRS, and TRH wrote the paper. All authors contributed intellectually to data analyses and interpretation.

## Acknowledgements

We thank Zuzana Storchová, Galal Metwalli, Marc Robinson-Rechavi, Gavin Conant, Camille Berthelot, and Jeremy Coate for valuable discussions.

