## Supplementary material for "Comparative regulomics reveals pervasive selection on gene dosage following whole genome duplication"

#### Gene ortholog groups

OrthoFinder was used to identify 30,818 gene ortholog groups (orthogroups) in seven fish species (Supplementary figure 1A) that comprised the majority of genes from each species (Supplementary figure 1C). After filtering by gene tree topology and minimum gene expression we identified 2,983 (10%) complete singleton orthogroups that contained only one gene copy from each species (referred to as singletons), and 4,053 (13%) complete duplicate orthogroups that contained only one gene copy from the non-salmonid species and two clades for the salmonids (Supplementary figure 1B). Orthogroups were still considered complete if they lacked a gene in zebrafish or medaka (but not both), but contained genes for all other species. We supplemented these complete orthogroups with 482 partial singleton groups and 2346 partial duplicate groups. Orthogroups were considered partial if they contained at least one salmonid gene in each salmonid clade, a pike gene and at least one zebrafish or medaka gene.

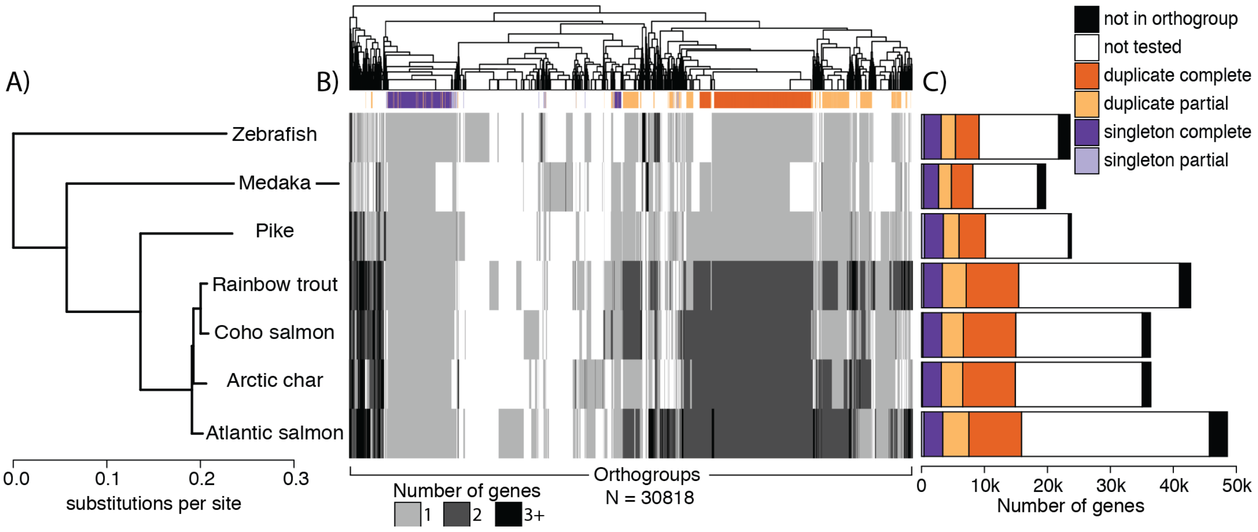


***Supplementary figure 1. Orthologs identified in the studied species.*** *(A) Phylogenetic tree of the seven species studied. (B) Heatmap showing an overview of the number of genes per species in each orthogroup. Complete singleton orthogroups have one gene copy in all salmonid species, and complete duplicate orthogroups have not more than one gene copy in the outgroup species and two gene copies (in separate clades) in all salmonid species. Partial groups lack some genes in one or more salmonid species. (C) Number of genes per species in orthogroups. Only complete/partial, singleton/duplicate orthogroups were tested for a significant shift in expression between the outgroups and the salmonids.*

#### Normalizing RNA-seq expression levels

Comparative analyses across species with widely different numbers of genes cannot be based on Transcripts Per Million (TPM) values directly. A naive expectation is that this will lead to lower gene expression values in species with higher numbers of genes, since the total number of reads is divided among more genes. However, this expectation is further complicated by the fact that the studied species also vary in the fraction of genes that are expressed in liver and the distribution of genes expressed at different levels (Supplementary figure 2A). To allow for comparison of gene expression across species, we therefore devised an approach where we first normalized expression data within each species (Supplementary figure 2B, between replicates) and then between species (Supplementary figure 2C). Between species normalization was done by computing scaling factors based on the expression of singletons, which we assumed to be expressed more similarly across species than duplicates. Species normalization factors differed quite substantially from the naive gene number expectation (NF in Supplementary figure 2C), with both within species and between species normalization having an effect on expression values. The normalized data behaves as expected with replicates clustering by species and species clustering in agreement with the phylogenetic species tree (Supplementary figure 3).

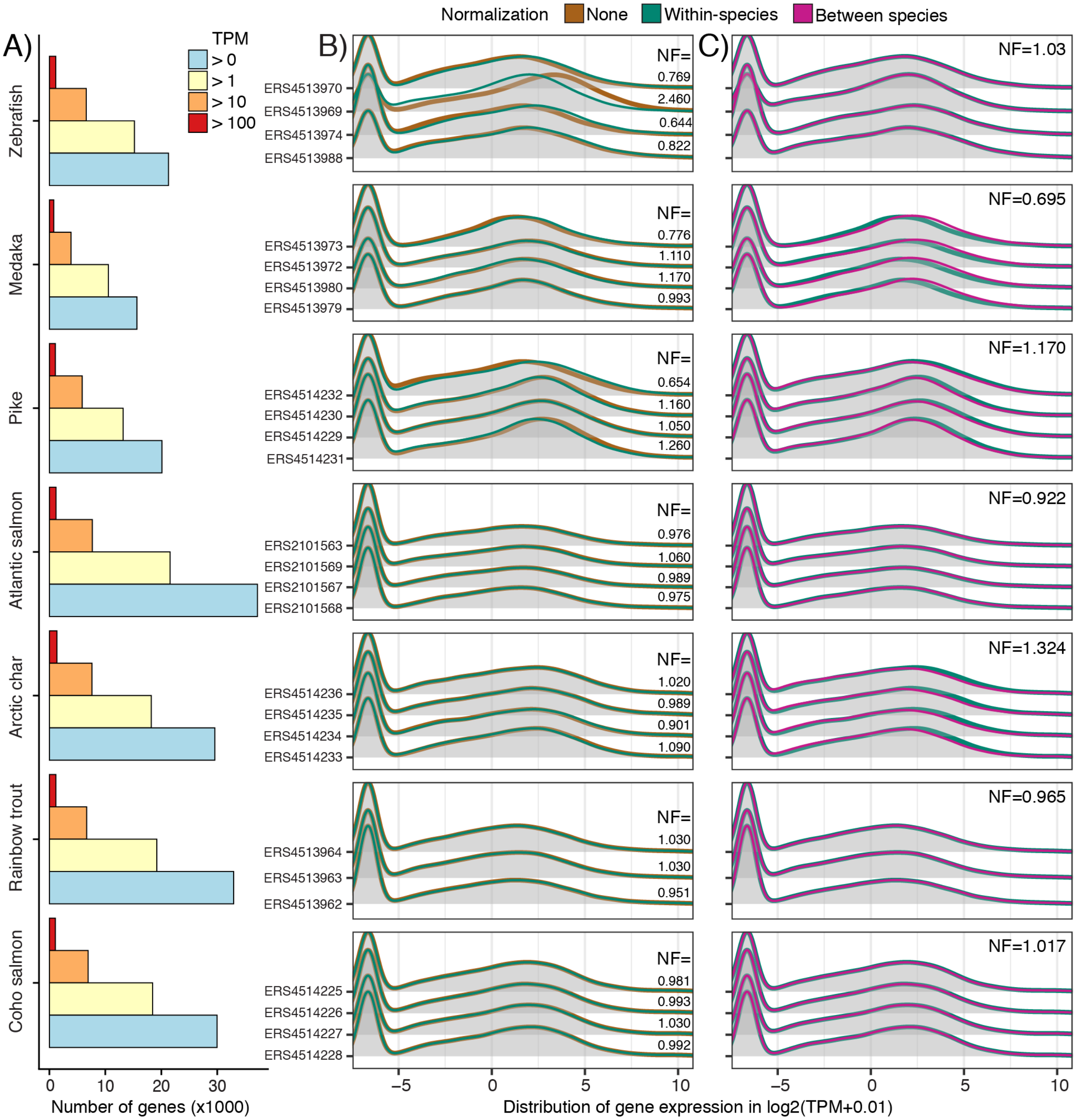


***Supplementary figure 2. Within and between species expression normalization.*** *(A) For each species, the number of genes that are expressed at different expression levels measured in Transcripts Per Million (TPM). (B) A comparison of the distribution of unnormalized gene expression values and within-species normalized values, for each replicate and species (sample names are IDs to ENA database entries). Values are log2 (TPM+0.01) and within-species normalization factors (NF) per sample is shown. The within-species normalization factor (NF) for each species is shown. (C) A comparison of the distribution of within-species gene expression values and between-species normalized values, for each replicate and species. The between-species normalization factor (NF) for each species is shown.*

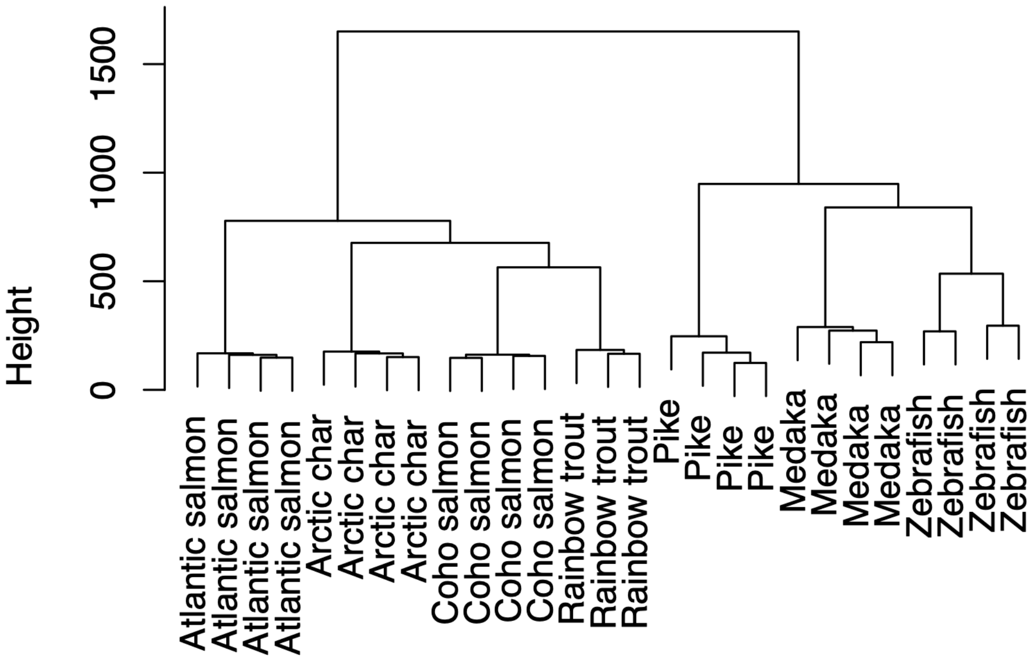


***Supplementary figure 3. Sample clustering.*** *Clustering of the 27 samples from the seven studied species. The tree displays the Euclidean distance between samples computed using the normalized expression levels from complete and partial ortholog groups. The tree was produced using the R function hclust with method = "ward.D".*

#### EVE method

The Ornstein-Uhlenbeck (OU) process has previously been used to model changes in gene expression over time (Chen et al. 2019; Brawand et al. 2011). The OU process is a stochastic process that models accumulation of random changes in expression over time (random walk), but unlike a Brownian motion process, an OU process assumes that expression level evolution is constrained within biologically relevant bounds (Supplementary figure 4A). Due to stabilizing selection, variation in expression increases non-linearly over time. In the OU process formulation, the $\theta$ parameter represents the optimal level that expression varies around. The $\sigma^2$ parameter represents the rate of variation over time, while the $\alpha$ parameter represents the strength of selection pressure towards an optimal level ($\theta$), restricting the variation (Supplementary figure 4A). To assess if expression variation in the orthologs from our studied species fits the OU process assumptions, we plotted the mean squared expression distance between pairs of species against their evolutionary distance (in sequence substitutions). Expression distance levels off as evolutionary distance increases (Supplementary figure 4B) in agreement with the OU process assumptions (rather than the Brownian motion process) (Bedford and Hartl 2009), demonstrating that an OU-model is appropriate for studying expression evolution in our species.

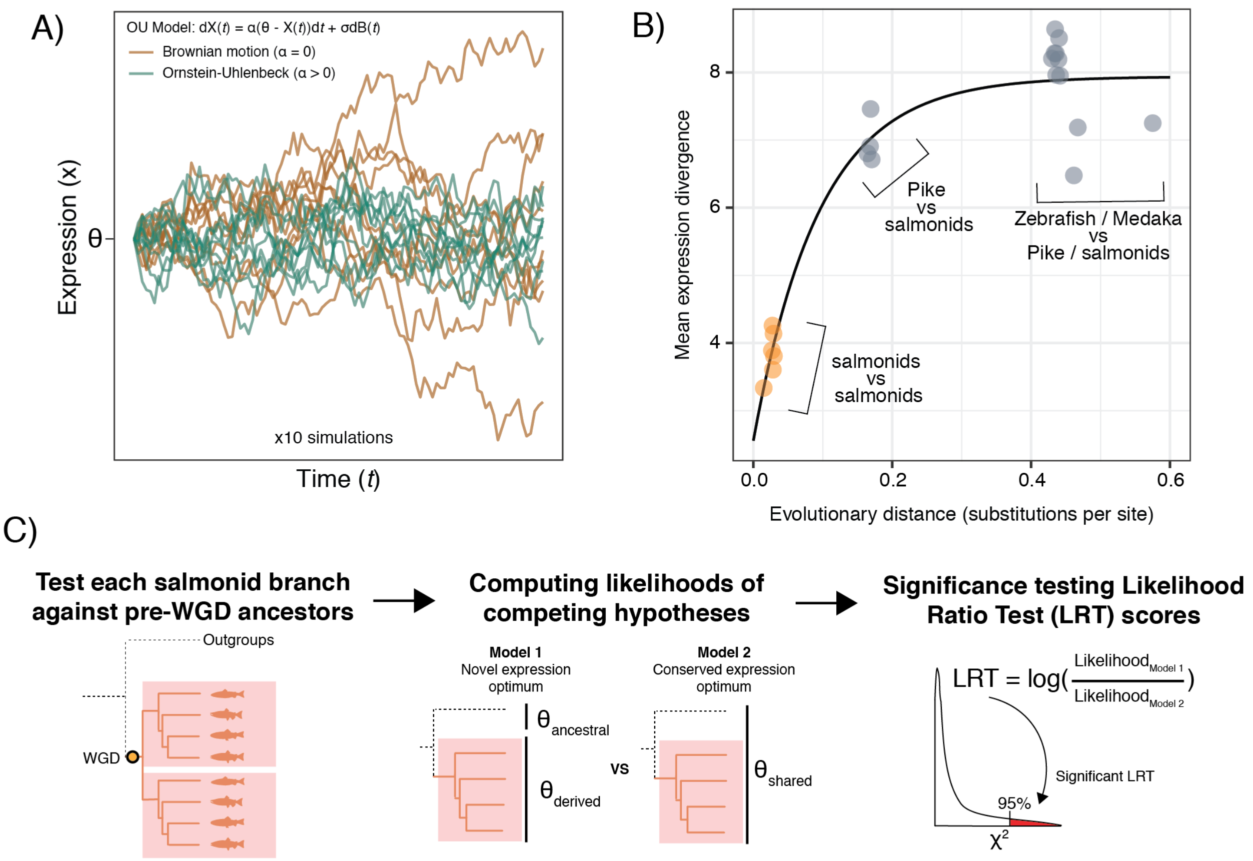


***Supplementary figure 4. Modeling gene expression divergence over time****. (A) Simulations of the variation in expression level over time comparing the Ornstein-Uhlenbeck (OU) process and the Brownian motion-based model. Parameters in the OU model include an optimal expression level θ, the rate of variation $\sigma^2$, and the strength of selection α towards the optimal level. (B) The mean squared expression distance is plotted against the evolutionary distance (sequence substitutions) for each pair of species in this study. The points are coloured and labeled based on the oldest species in the pair. (C) For each gene duplicate, we compare two models. Under model 1, the expression optimum of the duplicate ($\theta_{derived}$) is shifted compared to the ancestral expression optimum ($\theta_{ancestral}$), while under model 2 there is no shift in expression. A Likelihood Ratio Test (LRT) is then used to compare the two models and the statistical significance is determined using the* $\chi^2$ distribution with one degree of freedom.

To test for statistically significant shifts in expression after the salmonid-specific WGD (4R Whole Genome Duplication), we used the Expression Variance and Evolution (EVE) model (Rohlfs and Nielsen 2015; Rohlfs et al. 2014), which builds upon the OU process by adding a new parameter β that represents the ratio of within-species expression variance over between-species variance. For this paper we developed and implemented a user friendly version of EVE in R (https://gitlab.com/sandve-lab/evemodel).

The number of species (i.e. tips in gene trees) correlate with the statistical power to detect expression level shifts. This makes it difficult to directly compare the expression shift test statistics between retained Ss4R copies and genes that have returned to singleton copies. To overcome this obstacle, we subsetted each duplicate orthogroup, with retained Ss4R duplicates, performed the test separately on each of the two monophyletic ‘duplicate’ clades (Supplementary figure 4C). The outgroup orthologs (i.e. genes in the species without the Ss4R WGD) remain the same for both subsets. For each set of orthogroups, the EVE model parameters were optimised to best fit the expression data and maximum likelihood values were calculated. Our null hypothesis ($H_{null}$) is that expression has not shifted after WGD from a single optimal level ($\theta_{ancestral}$) and that expression differences are explained by species and evolutionary variance under the model of consistent stabilizing selection. Our alternative hypothesis ($H_{alt}$) is that expression has shifted for salmonid orthologs after WGD from a pre-Ss4R optimal level ($\theta_{ancestral}$) to a new post-Ss4R optimal level ($\theta_{derived}$). We then perform a Likelihood Ratio Test (LRT) and reject $H_{null}$ ($\theta_{ancestral} = \theta_{derived}$) and accept $H_{alt}$ ($\theta_{ancestral} \neq \theta_{derived}$) if the LRT score is greater than the upper 95% quantile of the $\chi^2$ distribution with one degree of freedom (Supplementary figure 4C). Salmonid genes with a significant shift in expression were further classified into those with a shift up and those with a shift down in expression compared to the pre-WGD level. The direction of the shift was determined by comparing the difference between $\theta_{derived}$ and $\theta_{ancestral}$. As controls, we used a corresponding setup to test for shifts from pre-Ss4R WGD levels in singleton orthogroups.

#### Expression level bias on expression shift detection

To test to what degree expression level biases our power to detect expression shifts in salmonids compared to the unduplicated outgroup species, we subdivided expression shift results (i.e the results shown in Figure 1C) into four quantiles based on the mean outgroup expression level (Supplementary figure 5). This analysis showed that the two main results are independent of expression level: there are more shifts in duplicated gene trees than in the singleton trees and the nature of these shifts are greatly skewed towards down-regulation in duplicated trees. However, there is a weaker trend showing that the power to detect shifts up increases with lower expression in the outgroups - for both duplicate and singleton trees. This is intuitive and unavoidable since there is little or no room for genes with low expression in the outgroup to be down-regulated in the salmonids, and vice versa for highly expressed genes.

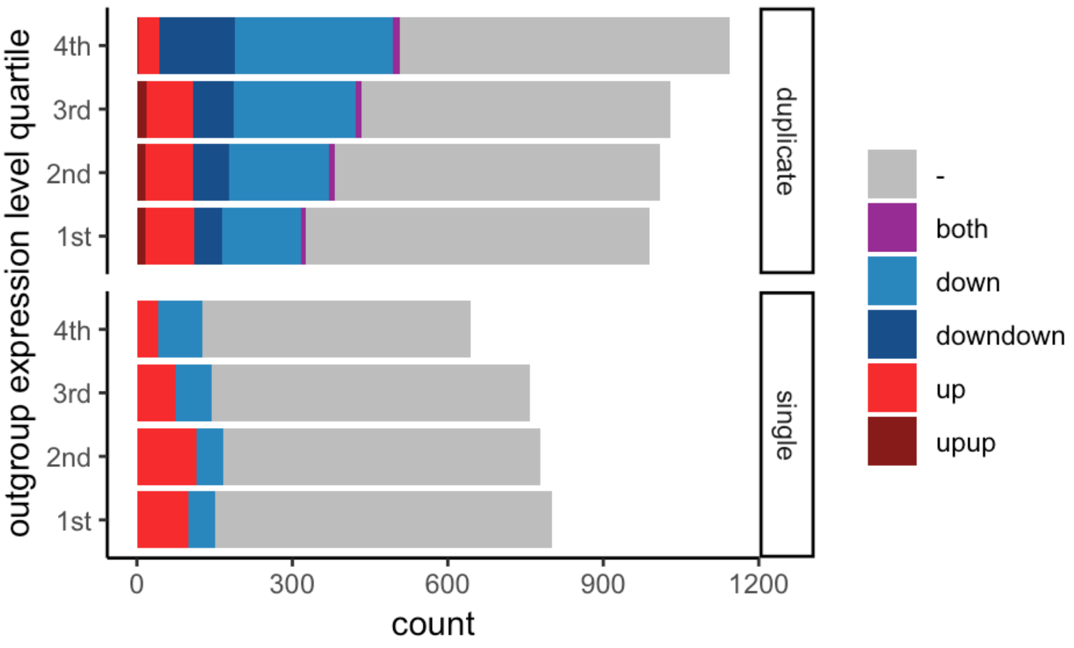

***Supplementary figure 5. Expression level bias on detecting expression shifts.*** *Expression shift results after dividing the tested gene trees (orthogroups) into four quantiles based on the mean outgroup expression, i.e. genes whose expression in the outgroup is ranked low / medium low / medium high / high.*

### Liver expression shifts in the context of a tissue atlas

We used an independent Atlantic salmon RNA-seq expression tissue atlas (Lien et al. 2016) to analyze complete and partial orthogroups with a significant expression shift (down or up) in salmonids compared to the outgroup species in one copy. These groups were chosen because the expression in the shifted copy can be compared to the conserved copy, which is assumed to retain ancestral expression levels. First, we confirmed that the copy with a liver expression shift detected by EVE also displays a shift in expression in the liver sample of the tissue atlas (Supplementary figure 6A). For up and down shifts, we then counted the number of tissues in the atlas that show a shift in expression between the conserved copy and the shifted copy in the same direction (Supplementary figure 6B). Partial groups show a higher proportion of ohnologs with a shift down in all 15 tissues compared to complete groups (Fisher's Exact Test, p = 3.062e-07). There was no corresponding significant difference between complete and partial groups for ohnologs with a shift up (p = 0.6484).

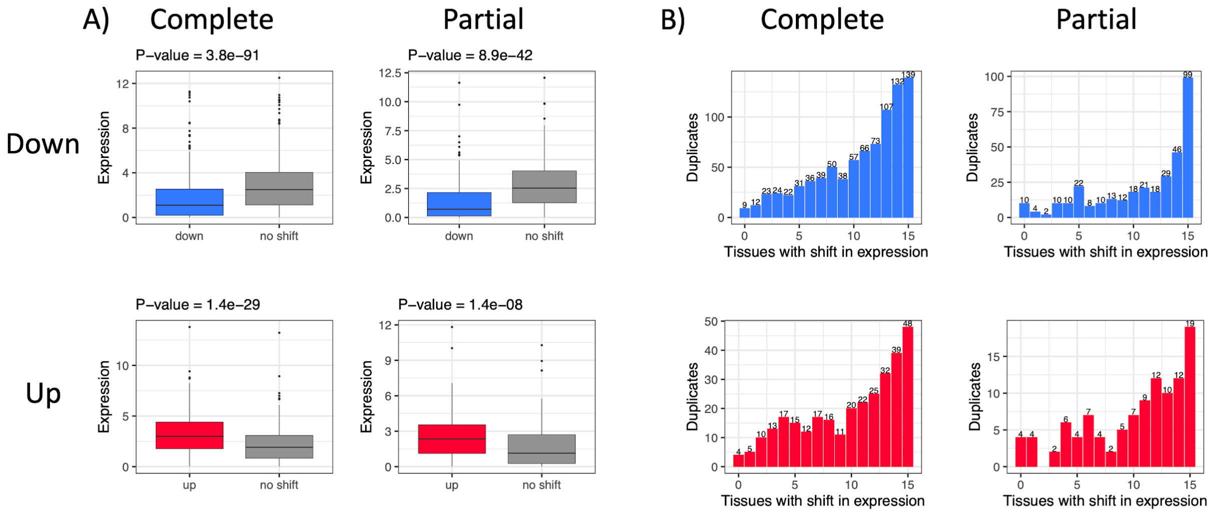


***Supplementary figure 6. Tissue expression atlas****. Analysis of detected expression shifts using an independent tissue atlas in Atlantic salmon containing 15 tissues. The plots are based on orthogroups with one copy having a significant expression shift down (blue) or up (red) in salmonids compared to the outgroup species - according to our EVE analysis. (A) Box plots compare the expression, in the liver sample of the tissue atlas, of the two copies, one shifted (down/up) and one remaining at ancestral levels (no shift). P-values were computed using the paired Wilcoxon test. (B) Bar plots of the number of tissues in the tissue altas with a shift up or down in the shifted copy compared to the copy without a shift.*

#### Selection pressure at the sequence level

We estimated branch specific selection pressure on coding sequences in the duplicate gene trees. Specifically, we calculated dN/dS measured at the branch from the WGD node to the root of each duplicate clade using the aBSREL method in Hyphy. The distribution of dN/dS values for each category of the expression shift results is shown in Supplementary figure 7. A two-sided paired wilcoxon test was performed to test if there is a difference in selection pressure between duplicates classified as differentially shifted (i.e. cons+down, cons+up, down+up). While the trend is that the lower expressed duplicate experience relaxed selection pressure, i.e. higher dN/dS, only the cons+down was significant at $p<0.05$, probably owing to the larger number of genes in this category.

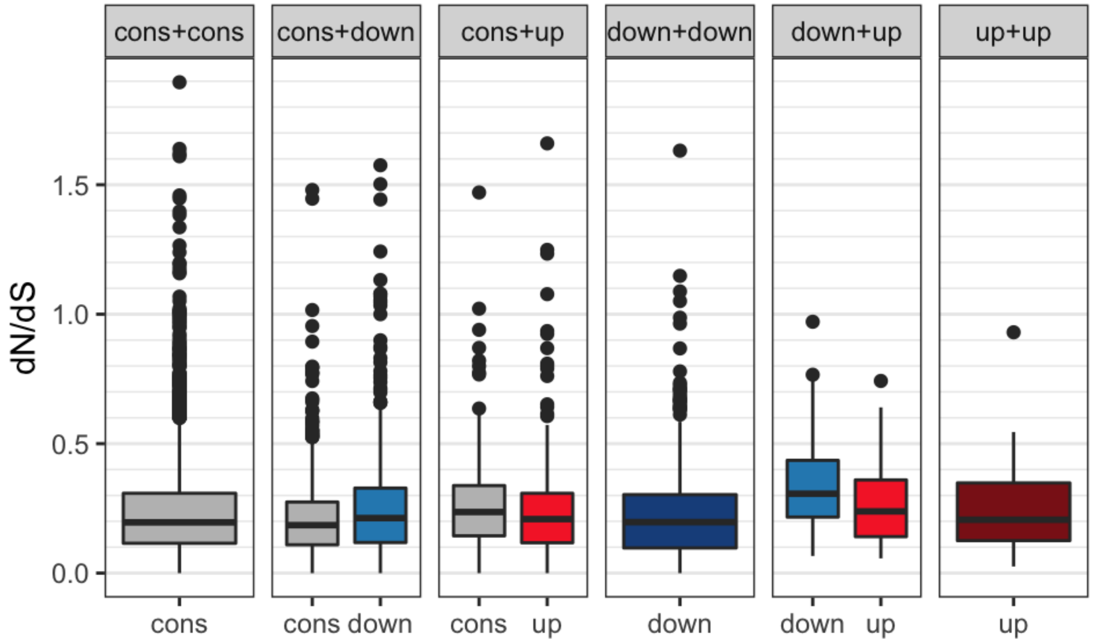


***Supplementary figure 7. dN/dS distribution per expression shift category.*** *A paired Wilcoxon test between duplicates with different EVE results within a category shows that only down vs cons have a significantly different dN/dS distribution.*

#### Transposable elements

For genes in the Atlantic salmon genome annotation (GCF_000233375.1), we defined gene promoter regions as 2000 bps upstream and 200 bps downstream of the gene's transcription start site (TSS). Transposable element (TE) annotations were taken from the Atlantic salmon genome paper (Lien et al. 2016). We calculated for each gene the proportion of promoter sequence that is overlapped by TE sequence using bedtools intersect of the promoter and TE annotations. We calculated overlap for all TE annotations, and separately for each TE superfamilies

(Supplementary figure 8). We compared the proportions of TE overlap between the conserved and shifted copy of ohnologs with an asymmetric down-shift in expression (Figure 1). We also compared a subset of these ohnologs that have an asymmetric down-shift in expression across all 15 tissues in an independently acquired tissue atlas (Supplementary figure 6). Ohnologs with promoters that did not overlap TEs for both copies were removed from the analysis. We tested the hypothesis that down-shift copies had a higher proportion of TE overlap than the conserved copy using a Wilcoxon paired test.

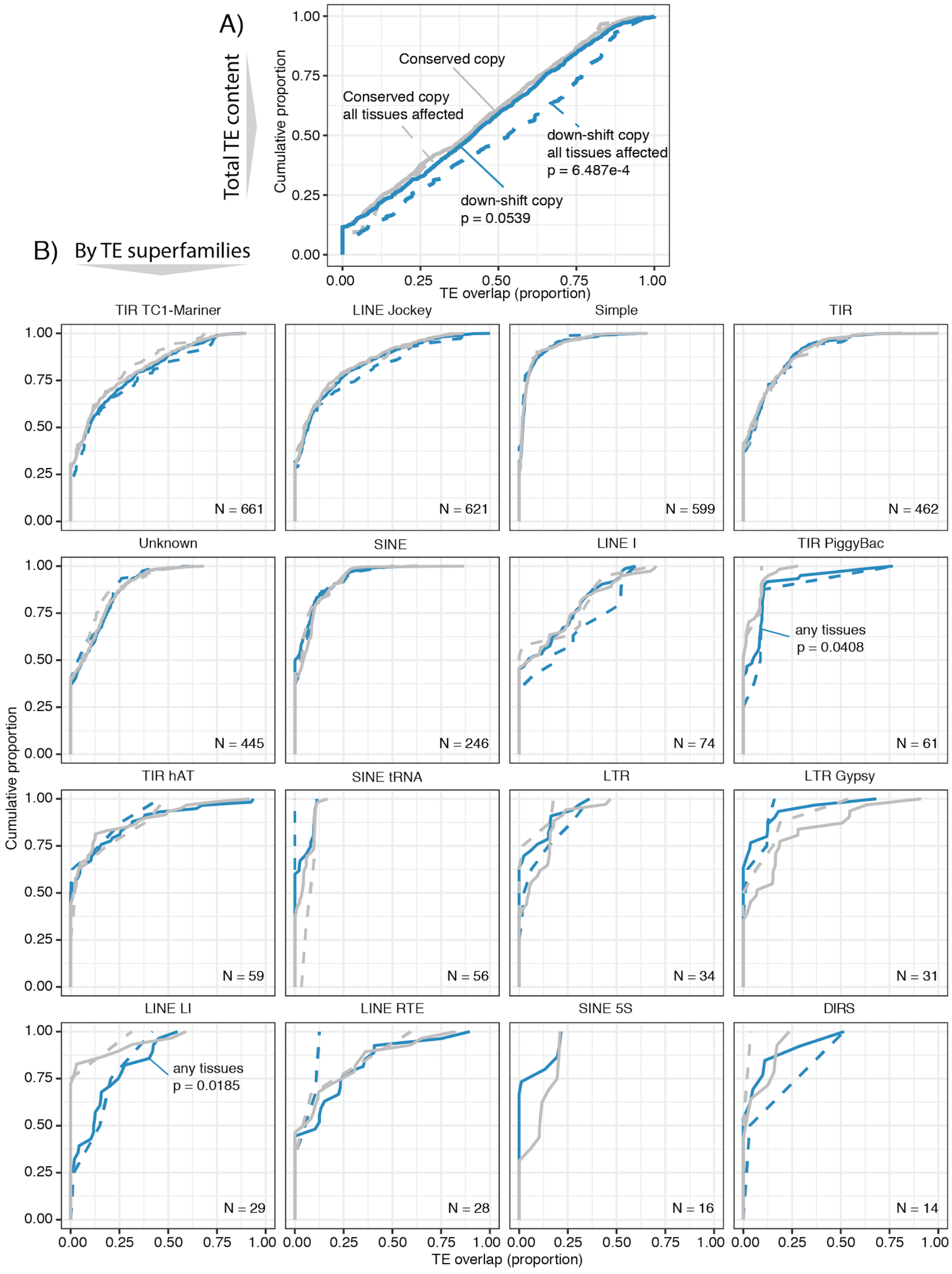

***Supplementary figure 8. Transposable elements in ohnolog promoters.*** *The amount of transposable element (TE) content in the promoters of Atlantic salmon ohnologs was measured as the proportion of the promoter sequence (2kb-/200+ of TSS) that is overlapped by sequence from (A) any TE, or (B) TE from a given superfamily classification (minimum 10 ohnologs). From the set of ohnologs with an asymmetric expression shift down, the cumulative proportions of TE overlap are shown for the conserved (grey) and down-shift (blue) copy. Overlaps are also shown for a subset of ohnologs with a down-shift across all (15) tissues in a tissue atlas (dashed lines). The hypothesis that down-shift copies had more TE overlap than conserved copies were tested using a Wilcoxon paired test (significant p-values are shown).*

### Fractionation bias

Large-scale differences in selective constraints on different copies of a duplicated genome can lead to biased fractionation (i.e. differences in ohnolog loss rate across duplicated regions). This has been observed in other fish genomes following the WGD in a teleost ancestor and in the allopolyploid common carp (Conant 2019; Xu et al. 2019). To test for such large-scale bias in purifying selection pressure after the salmonid WGD, we quantified biases in ohnolog evolution between large conserved duplicated syntenic regions (Supplementary figure 9A). Nine out of 47 syntenic duplicated blocks (covering 80% of the genome) had significantly biased duplicate gene loss rates (i.e. gene fractionation) (Supplementary figure 9B), but we did not find significant bias in expression fractionation, defined as loss of expression level in one ohnolog copy (Supplementary figure 9C). Nevertheless, we found a weak but significant correlation between gene and expression fractionation (Supplementary figure 9C). This suggests that the test power for expression fractionation for each duplicated region is low, but that gene and expression fractionation processes could be mechanistically linked. In conclusion we find that asymmetric relaxation of purifying selection pressure on one ohnolog copy is a major driver of regulatory evolution following WGD, but that this asymmetry is only weakly linked to bias in selective constraints across larger duplicated genomic regions.

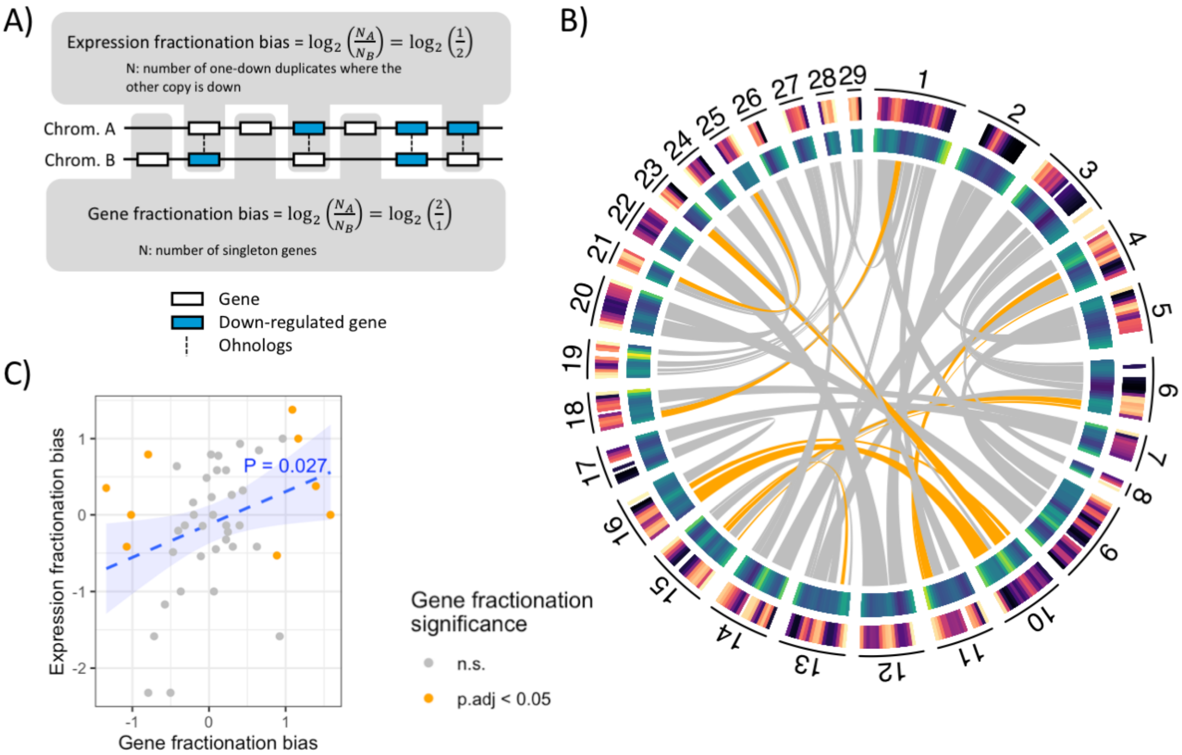


***Supplementary figure 9. Fractionation bias.*** *(A) The figure illustrates genes aligned along a pair of ohnologuous chromosome regions. Expression fractionation bias represents the tendency for genes on one of the chromosomes being expressed at a generally higher level than their corresponding duplicates on the other chromosome. It is calculated by considering the duplicate pairs where one of the duplicates is down-shifted, and taking the ratio between the number of non-shifted genes in each chromosome. Gene fractionation bias is calculated by considering the singleton genes, i.e. genes whose duplicates have been lost, and taking the ratio of the number of retained genes in each of the duplicated chromosome regions. (B) Circos plot showing syntenic duplicated blocks, i.e. ohnologuous chromosome regions (links). Orange coloured links indicate significant gene fractionation bias. Outer and inner circular heatmap corresponds to expression and gene fractionation, respectively (lighter colour = more fractionation). (C) Gene and expression fractionation ratio per duplicated synteny block. Gene fractionation ratio is the ratio of the number of retained singletons in each of the duplicated regions. E.g. if there are twice as many lost in one region compared to the duplicate region, the log2 ratio is 1 or -1 (order is arbitrary). Expression fractionation ratio is the ratio of the number of retained duplicates where one copy is not down while the other copy is down-regulated in each of the duplicated regions (see panel (A)). Orange colored dots indicate significant bias of gene fractionation (binomial test, FDR < 0.05). There was no significant bias of expression fractionation.*

### Genes function enrichment - up/down shift

***Supplementary table 1. KEGG pathway enrichment for genes with expression shifts up or down.***

| Expression shift | Pathway | N in pathway | N with shift | P-value |
|---|---|---|---|---|
| up | Fatty acid elongation | 12 | 4 | 2.50E-02 |
| up | Fatty acid metabolism | 24 | 6 | 2.67E-02 |
| up | Cell cycle | 61 | 11 | 3.60E-02 |
| down | Ribosome | 43 | 34 | 5.59E-11 |
| down | Oxidative phosphorylation | 53 | 26 | 3.29E-03 |
| down | Ribosome biogenesis in eukaryotes | 28 | 15 | 8.79E-03 |
| down | NOD-like receptor signaling pathway | 73 | 31 | 1.90E-02 |
| down | Phagosome | 55 | 24 | 2.57E-02 |
| down | FoxO signaling pathway | 83 | 34 | 2.59E-02 |
| down | Aminoacyl-tRNA biosynthesis | 9 | 6 | 2.70E-02 |
| down | Sphingolipid metabolism | 21 | 11 | 2.90E-02 |
| down | Toll-like receptor signaling pathway | 42 | 19 | 3.00E-02 |

***Supplementary table 2. Cell cycle genes with potential roles in genomic stability identified among the up-regulated ohnologs.*** *These genes were identified either via KEGG analyses done on all up-regulated genes (Supplementary table 1) or separately for each expression shift category (Supplementary table 4) and are supplemented by manual identifications (NA in the column 'KEGG enrichment'). The inference of orthology with human and yeast cell cycle proteins is described in Supplementary Figure 10.*

| Gene ID | Product | KEGG ID | Pathway | KEGG enrichment | Shift category | Human Ortholog | Human % ID | Yeast Ortholog | Yeast % ID |
|---|---|---|---|---|---|---|---|---|---|
| 106563208 | 14-3-3 protein eta-like | 04110 | Cell cycle | 'all up' 'up' | 'up' | YWHAH | 84 | BMH2 | 58 |
| 106569538 | Fizzy-related protein homolog | 04110 | Cell cycle | 'all up' 'up' | 'up' | FZR1 | 90 | CDH1 | 46 |
| 106569474 | Zygotic DNA replication licensing factor mcm6-B-like | 04110 | Cell cycle | 'all up' 'up' | 'up' | MCM6 | 80 | MCM6 | 43 |
| 106565437 | Cyclin-dependent kinase 2-like | 04115 04110 | Cell cycle; p53 signaling pathway | 'all up' 'up' | 'up' | CDK2 | 91 | CDC28 | 62 |
| 106585187 | Dual specificity protein phosphatase CDC14A-like | 04110 | Cell cycle | 'all up' 'up' | 'up' | CDC14A | 63 | - | N.A |
| 106572360 | Transcription factor E2F1-like | 04110 | Cell cycle | 'all up' 'up' | 'up' | E2F1 | 41 | CDC14 | 31 |
| 106587338 | G2/mitotic-specific cyclin-B2 | 04115 04110 | Cell cycle; p53 signaling pathway | 'all up' 'up' | 'up' | CCNB2 | 58 | CLB4 | 38 |
| 100196122 | Cell division cycle 2 | 04115 04110 | Cell cycle; p53 signaling pathway | 'all up' 'up' | 'up' | CDK1 | 86 | CDC28 | 61 |
| 106583830 | Dual specificity protein kinase Ttk-like | 04110 | Cell cycle | 'all up' 'up' | 'up' | TTK | 40 | MPS1 | 33 |
| 106588946 | Cyclin-dependent kinase 6-like | 04115 04110 | Cell cycle; p53 signaling pathway | 'all up' 'up' | 'up' | CDK6 | 82 | CDC28 | 48 |
| 106569074 | E3 ubiquitin-protein ligase RFWD2-like | 04115 | p53 signaling pathway | 'up' | 'up' | RFWD2 | 85 | - | - |
| 106562012 | G1/S-specific cyclin-E1-like | 04115 | p53 signaling pathway | 'up' | 'up up' | CCNE1 | 63 | CLB5 | 37 |
| 100194938 | X-ray repair cross-complementing protein 6 | 03450 | Non-homologous end-joining | 'up' | 'up' | XRCC6 | 54 | - | - |
| 100196000 | flap endonuclease 1 | 03450 | Non-homologous end-joining | 'up' | 'up' | FEN1 | 74 | RAD27 | 55 |
| 100380415 | cyclin-Y isoform X3 | NA | NA | NA | 'up' | CCNY | 89 | N.A | N.A |
| 106565771 | host cell factor 1-like | NA | NA | NA | 'up' | HCFC1 | 62 | - | - |
| 106580490 | cell division cycle-associated protein 2-like | NA | NA | NA | 'up' | - | - | - | - |
| 106607947 | cyclin-C | NA | NA | NA | 'up' | CCNC | 96 | SSN8 | 28 |
| 106573334 | PCNA-associated factor-like | NA | NA | NA | 'up' | KIAA0101 | 63 | - | - |
| 100380429 | regulator of chromosome condensation | NA | NA | NA | 'up' | RCC1 | 74 | SRM1 | 30 |
| 106604944 | casein kinase I isoform alpha | NA | NA | NA | 'up' | CSNK1A1 | 99 | HRR25 | 61 |
| 106571202 | denticleless protein homolog | NA | NA | NA | 'up' | DTL | 49 | N.A | N.A |
| 106612241 | cyclin-J-like | NA | NA | NA | 'up up' | CCNJ | 49 | N.A | N.A |
| 106563510 | chromosome-associated kinesin KIF4-like | NA | NA | NA | 'up up' | KIF4B | 64 | KIP1 | 36 |
| 106562237 | uncharacterized protein LOC106562237 | NA | NA | NA | 'up up' | CENPT | 37 | - | - |
| 106588468 | claspin-like | NA | NA | NA | 'up up' | CLSPN | 53 | - | - |
| **106563621** | kinesin-like protein KIF20A | NA | NA | NA | 'up up' | KIF20A | 47 | KIP2 | 23 |
| **106612533** | microcephalin | NA | NA | NA | 'up up' | MCPH1 | 35 | - | - |
| **106597351** | serine/threonine-protein kinase PLK4-like | NA | NA | NA | 'up up' | PLK4 | 52 | CDC5 | 39 |

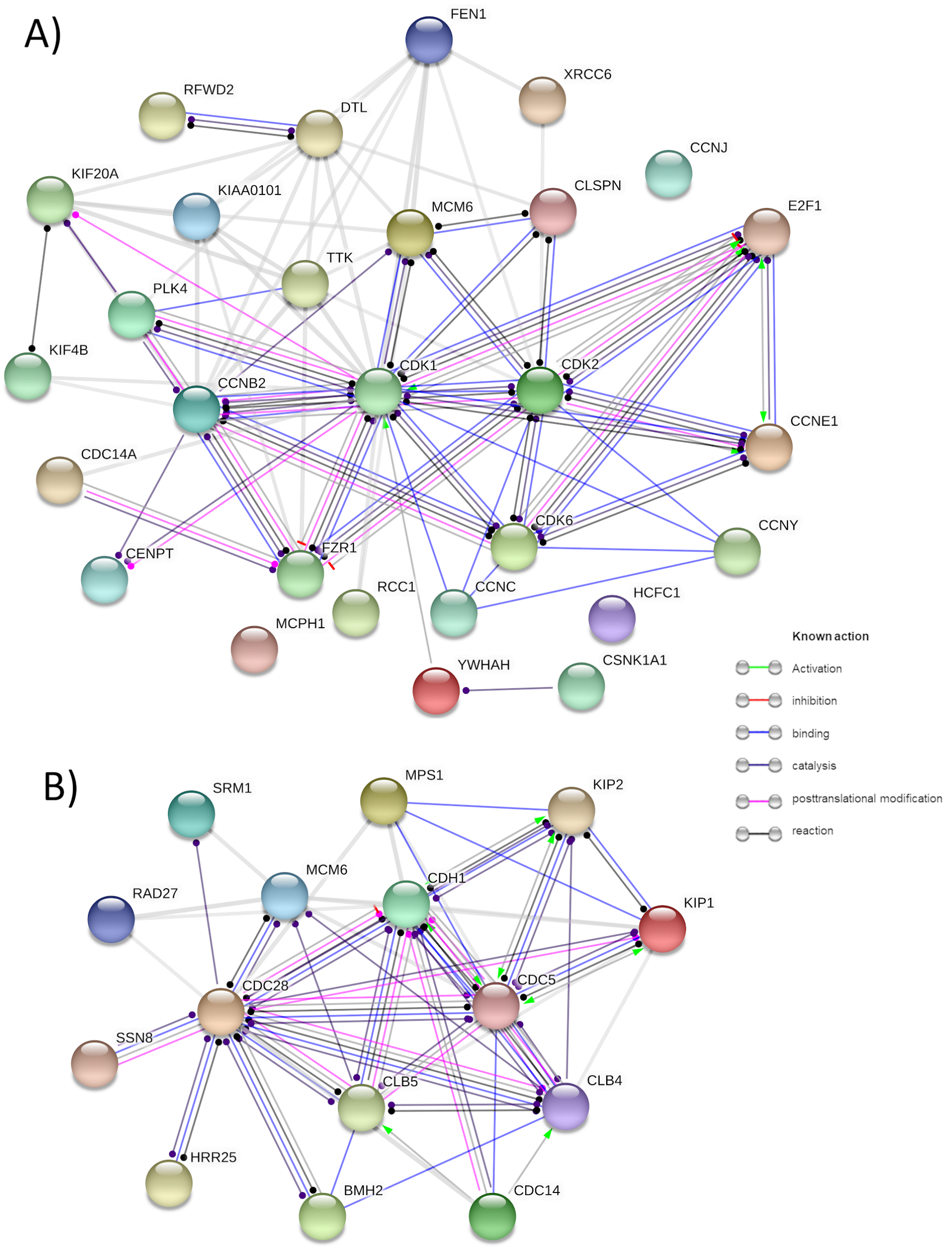


***Supplementary figure 10. Upshifted ohnologs include many highly conserved genes encoding proteins controlling cell division.*** *Protein-protein interaction (PPI) network visualizations are shown for human (A) and yeast (B) orthologs of the products of 29 salmonid genes shown in Supplementary Table 2. These visualizations and the inference of orthology were performed in STRING (Szklarczyk et al. 2015) using a minimum PPI confidence score of 0.7 ('high confidence'). For human orthologs, 10 PPIs were expected when 76 are observed (p < 1.0e-16). For yeast orthologs, 5 PPIs were expected when 47 are observed (p < 1.0e-16). This network centres on cyclin proteins (Cyclin B, C, E, J and Y) and cyclin-dependent kinases (Cdk1, Cdk2, Cdk6) that form complexes required for different phases of mitosis (Gjelsvik et al. 2019), along with proteins governing replication (Mcm6), cell cycle progression (E2f1), anaphase (Fzr1), centrosome separation (Cdc14a), cell cycle arrest (Claspin), centriole duplication (Plk4), cytokinesis (KIF20A) and chromosome segregation (Cenpt), condensation (Rcc1) and alignment (Ttk). In many cases, the up-shift was not tissue-specific, e.g. up-regulation of one ohnolog encoding Cdk1, which shares >60% identity between salmonids and yeast, was present in all 15 tissues. These results indicate that conserved functional pathways involved in the regulation of genomic stability were targeted for increased dosage after WGD.*

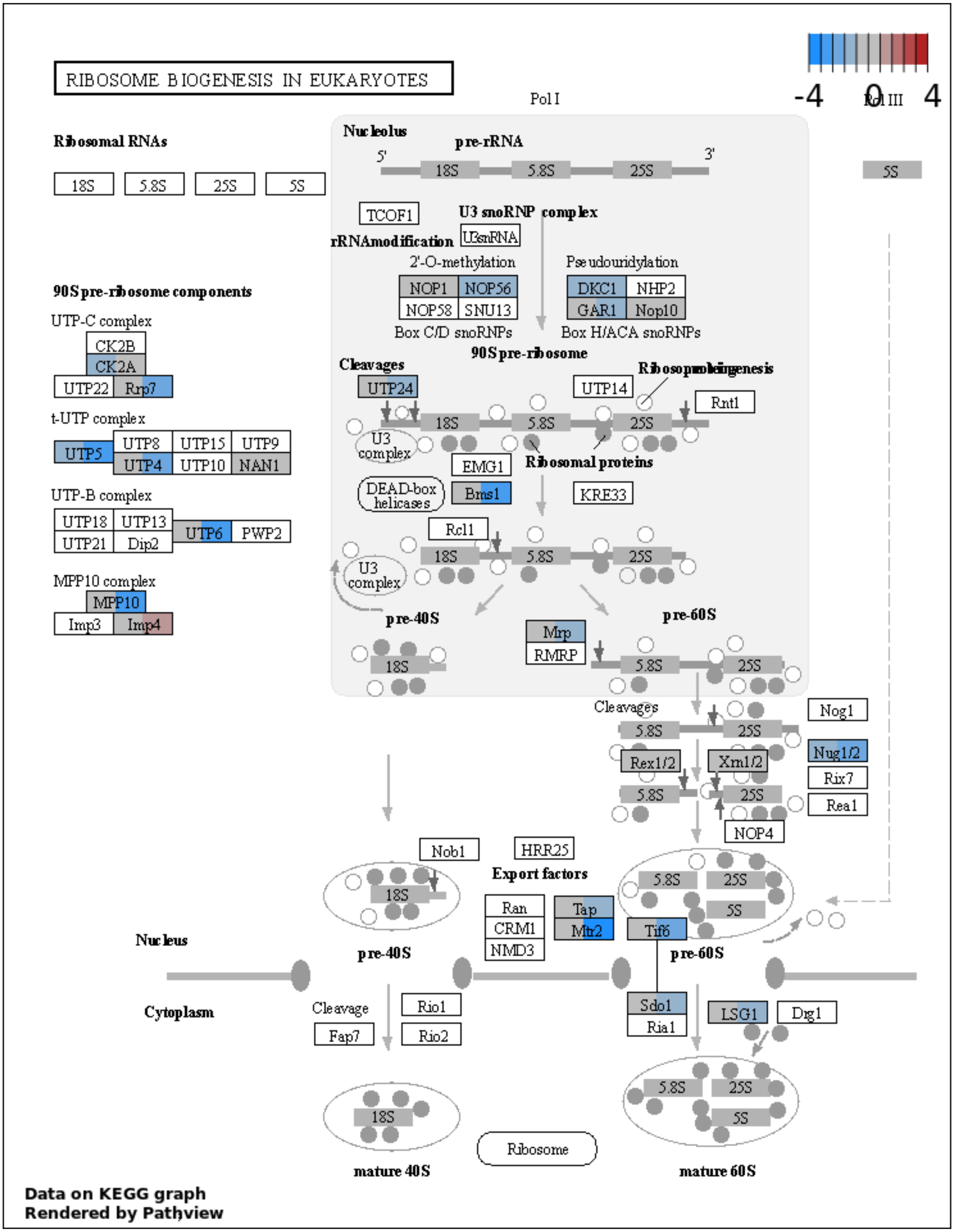


***Supplementary figure 11. Ohnolog expression in ribosome biogenesis pathway.*** *Each gene node (rectangle) displays two expression values for duplicate 1 and duplicate 2 (order based on highest-lowest shift p-value). These*

*values are the difference in the mean log expression values for the salmonids minus the outgroups mean log expression. For multiple genes at a node the mean difference is calculated.*

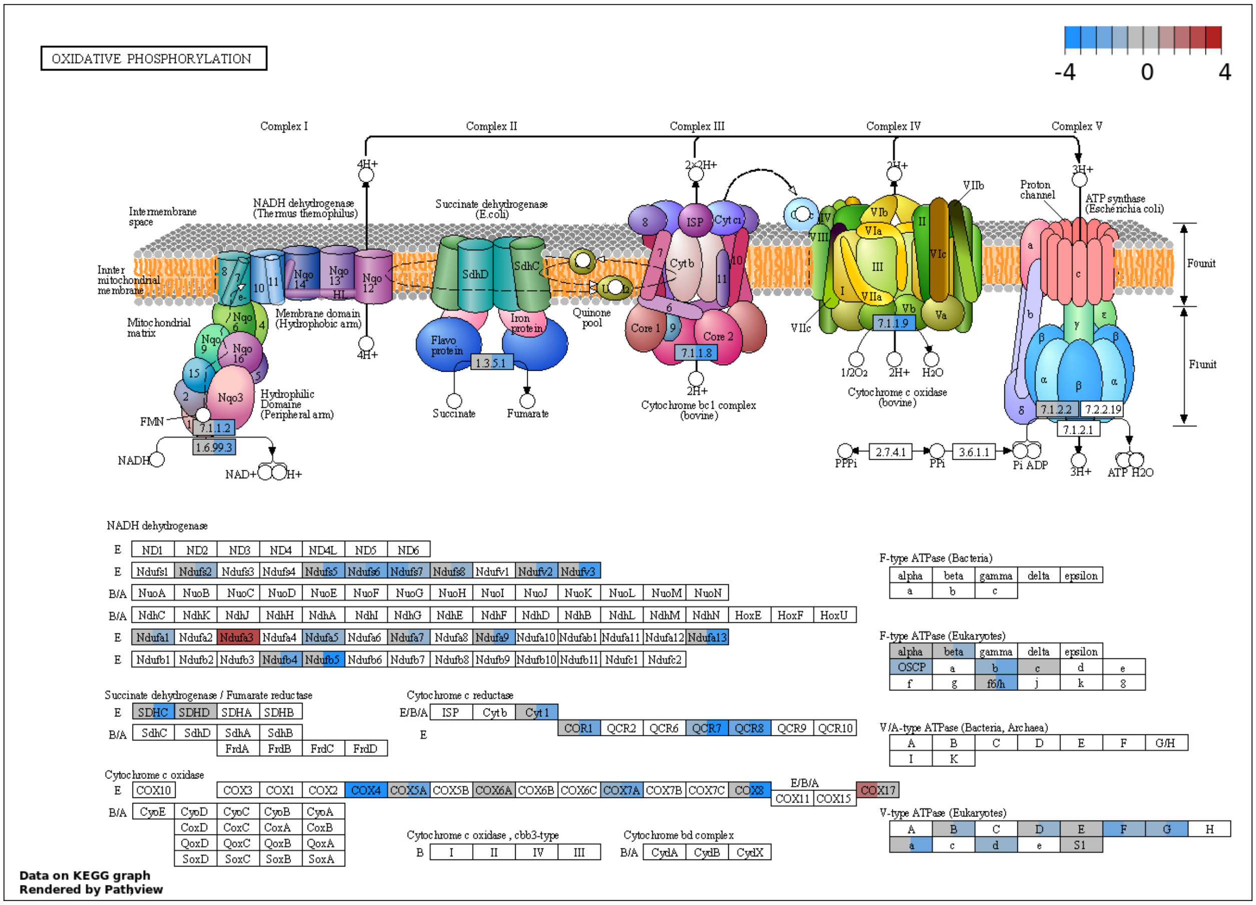


***Supplementary figure 12. Ohnolog expression in oxidative phosphorylation pathway.*** *Each gene node (rectangle) displays two expression values for duplicate 1 and duplicate 2 (order based on highest-lowest shift p-value). These values are the difference in the mean log expression values for the salmonids minus the outgroups mean log expression. For multiple genes at a node the mean difference is calculated.*

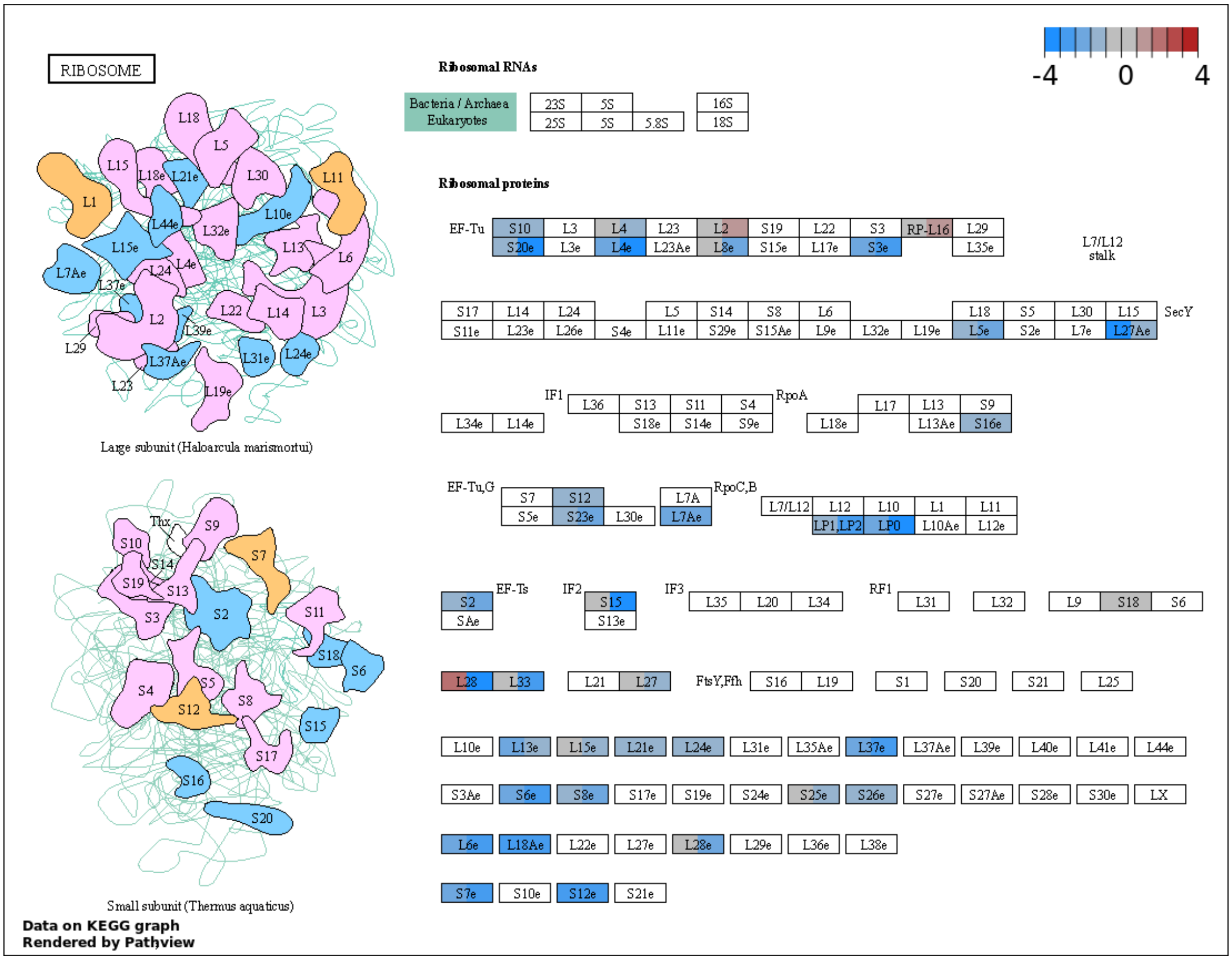


***Supplementary figure 13. Ohnolog expression in ribosome pathway.*** *Each gene node (rectangle) displays two expression values for duplicate 1 and duplicate 2 (order based on highest-lowest shift p-value). These values are the difference in the mean log expression values for the salmonids minus the outgroups mean log expression. For multiple genes at a node the mean difference is calculated.*

### Protein complexes

We assigned orthogroups as being in a protein complex or not based on the human ortholog and it's protein complex annotations from the CORUM database (Giurgiu et al. 2019). We used a Fisher's exact test, for singleton and ohnolog genes in an expression shift category (see legend in Figure 1), to test whether more genes within an expression shift category were in a protein complex than expected by chance (Supplementary table 3, Supplementary figure 14). We tested using the set of all complexes and a subset of complexes that contained a mix of both singtons and ohnologs.

***Supplementary table 3. Enrichment of shifted orthogroups in protein complexes.*** *We tested if orthogroups within each shift category (see legend Figure 1) were enriched in protein complexes - for singleton and ohnolog orthogroups. P-values were calculated using the Fisher's exact test. We tested using all complexes, and the set of complexes that contained a mix of both singletons and ohnologs.*

| Complex type | Orthogroup type | Expression shift category | N total in complex | N total not in complex | N shifted in complex | N shifted not in complex | P-value |
|---|---|---|---|---|---|---|---|
| all | singleton | up | 633 | 2832 | 80 | 277 | 0.0348 |
| all | singleton | down | 633 | 2832 | 57 | 280 | 0.7527 |
| all | ohnolog | up | 1463 | 5226 | 119 | 382 | 0.1773 |
| all | ohnolog | up + up | 1463 | 5226 | 15 | 76 | 0.9176 |
| all | ohnolog | down | 1463 | 5226 | 315 | 1106 | 0.4155 |
| all | ohnolog | down + down | 1463 | 5226 | 106 | 435 | 0.9027 |
| all | ohnolog | both | 1463 | 5226 | 19 | 54 | 0.2345 |
| single and ohnolog | singleton | up | 532 | 2933 | 66 | 291 | 0.0721 |
| single and ohnolog | singleton | down | 532 | 2933 | 49 | 288 | 0.6785 |
| single and ohnolog | ohnolog | up | 983 | 5706 | 75 | 426 | 0.4543 |
| single and ohnolog | ohnolog | up + up | 983 | 5706 | 13 | 78 | 0.5894 |
| single and ohnolog | ohnolog | down | 983 | 5706 | 210 | 1211 | 0.4821 |
| single and ohnolog | ohnolog | down + down | 983 | 5706 | 75 | 466 | 0.7197 |
| single and ohnolog | ohnolog | both | 983 | 5706 | 15 | 58 | 0.111 |

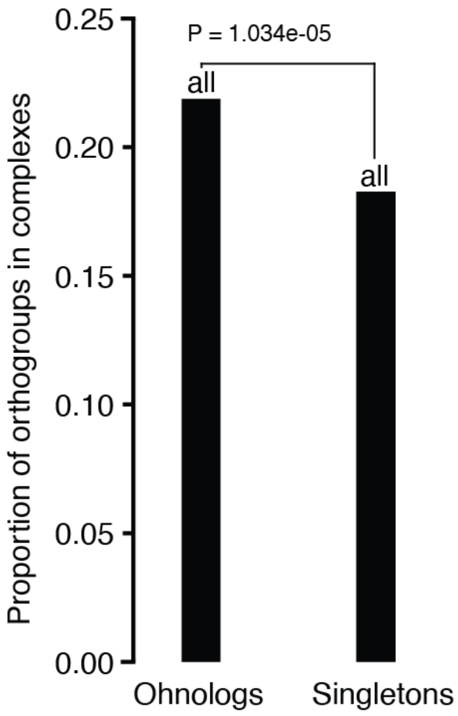


***Supplementary figure 14. Protein complexes****. Fraction of ohnologs versus singletons in protein complexes. One-sided Fisher's exact test test p = 1e-5.*

#### Gene function enrichment - evolutionary categories

We assigned KEGG pathway annotations to the orthogroups based on the Northern pike ortholog and it's KEGG annotations. We then tested each set of ohnologs within an expression shift category for the enrichment of KEGG pathways using the kegga function from the limma R package, with the total set of tested ohnologs as the background.

***Supplementary table 4. KEGG pathway enrichment for sets of ohnologs with expression shifts.***

| Expression shift category | KEGG ID | Pathway | N ohnologs in pathway | N ohnologs in category | P-value |
|---|---|---|---|---|---|
| opposite | 00563 | Glycosylphosphatidylinositol (GPI)-anchor biosynthesis | 7 | 2 | 2.38E-03 |
| opposite | 00062 | Fatty acid elongation | 12 | 2 | 7.22E-03 |
| opposite | 04530 | Tight junction | 89 | 4 | 1.59E-02 |
| opposite | 04514 | Cell adhesion molecules (CAMs) | 61 | 3 | 2.86E-02 |
| opposite | 01212 | Fatty acid metabolism | 25 | 2 | 3.00E-02 |
| opposite & down | 01100 | Metabolic pathways | 583 | 13 & 145 | 9.58E-03 & 1.54E-02 |
| down | 00190 | Oxidative phosphorylation | 53 | 24 | 7.31E-05 |
| down | 03008 | Ribosome biogenesis in eukaryotes | 28 | 14 | 6.94E-04 |
| down | 04621 | NOD-like receptor signaling pathway | 71 | 25 | 4.51E-03 |
| down | 04146 | Peroxisome | 24 | 10 | 1.89E-02 |
| down | 04142 | Lysosome | 51 | 17 | 3.03E-02 |
| down | 04130 | SNARE interactions in vesicular transport | 16 | 7 | 3.61E-02 |
| down | 00630 | Glyoxylate and dicarboxylate metabolism | 13 | 6 | 3.94E-02 |
| down | 00350 | Tyrosine metabolism | 10 | 5 | 4.16E-02 |
| down + down | 03010 | Ribosome | 43 | 20 | 1.68E-11 |
| down + down | 04672 | Intestinal immune network for IgA production | 10 | 4 | 5.99E-03 |
| down + down | 04371 | Apelin signaling pathway | 81 | 12 | 2.86E-02 |
| down + down | 00970 | Aminoacyl-tRNA biosynthesis | 9 | 3 | 3.06E-02 |
| down + down | 00471 | D-Glutamine and D-glutamate metabolism | 4 | 2 | 3.51E-02 |
| down + down | 04623 | Cytosolic DNA-sensing pathway | 16 | 4 | 3.52E-02 |
| down + down | 04620 | Toll-like receptor signaling pathway | 40 | 7 | 3.91E-02 |
| up | 04110 | Cell cycle | 60 | 10 | 1.28E-02 |
| up | 03450 | Non-homologous end-joining | 4 | 2 | 3.03E-02 |
| up | 04115 | p53 signaling pathway | 35 | 6 | 4.33E-02 |
| up + up | 00524 | Neomycin, kanamycin and gentamicin biosynthesis | 2 | 1 | 2.70E-02 |
| up + up | 00270 | Cysteine and methionine metabolism | 24 | 2 | 4.16E-02 |

### ATAC-Seq data validation

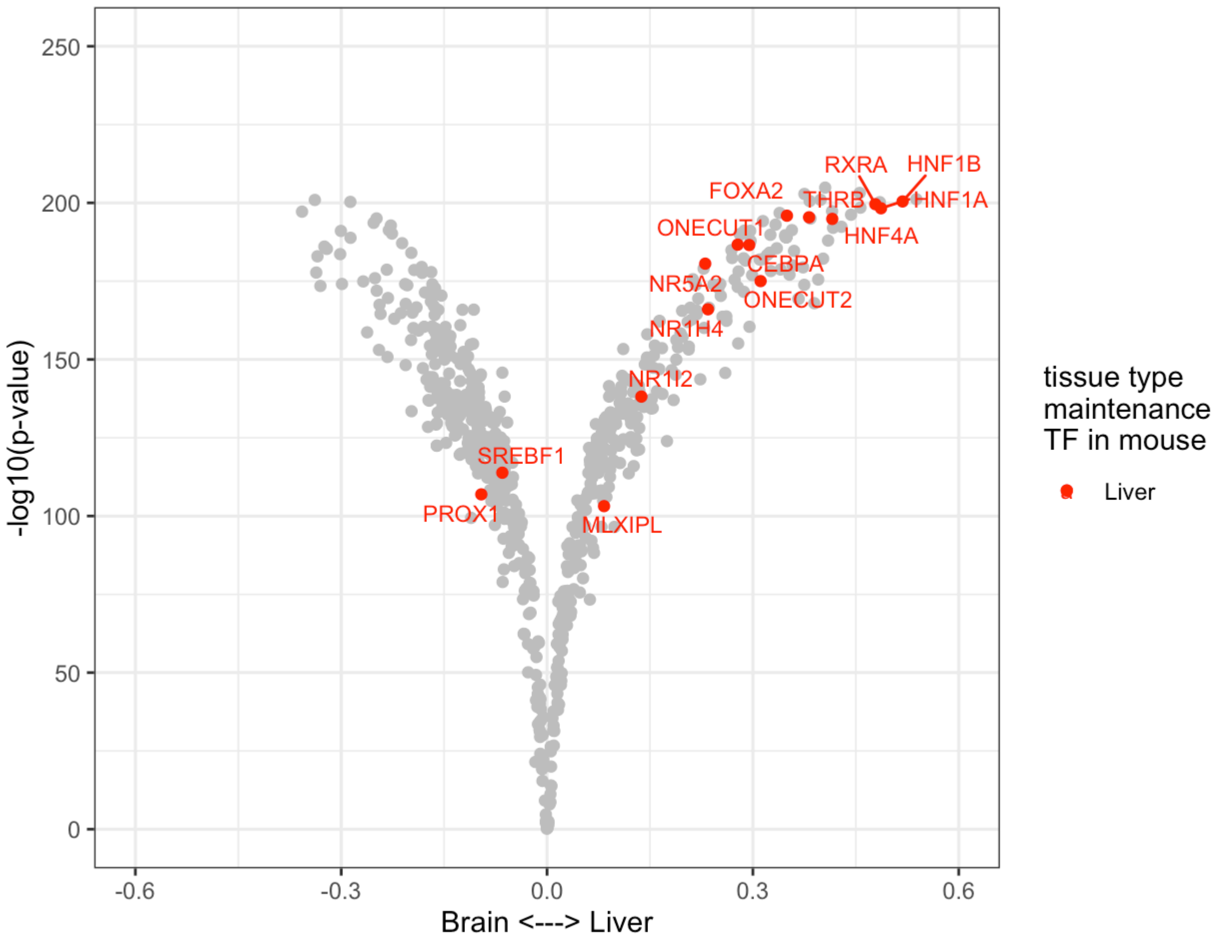


***Supplementary figure 15. Comparison of TF activity between Liver and Brain**. Volcano plot showing the TF binding activity difference between brain and liver in Atlantic salmon estimated from ATAC-seq using TOBIAS. Each dot represents a motif from the JASPAR core vertebrate TF database. The labeled motifs refer to liver tissue type maintenance TFs, from a study on mouse (Zhou et al. 2017).*

### Bound TFBS analysis

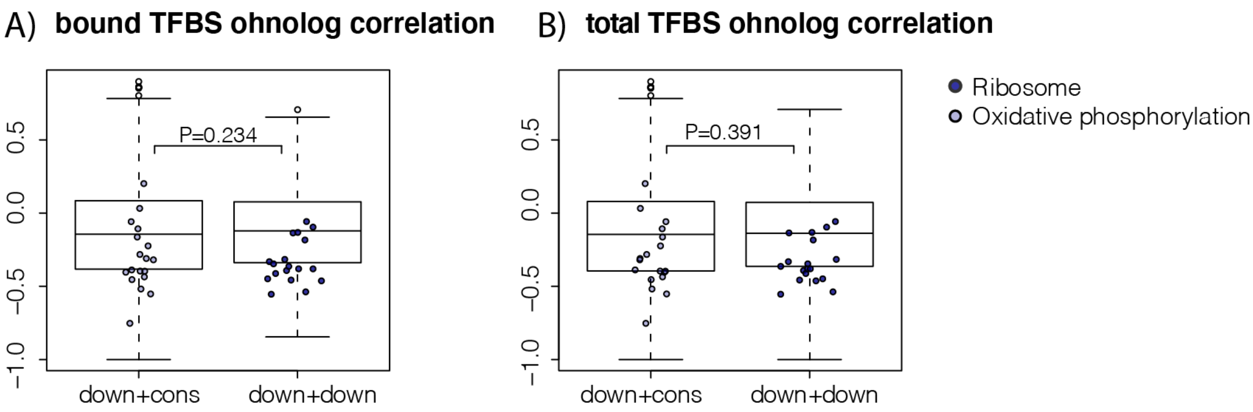


***Supplementary figure 16: Similarity of bTFBSs in promoters**. Comparison of (A) bTFBSs and (B) TFBSs in promoters of asymmetrically evolving (down+cons) and symmetrically evolving (down+down) ohnolog pairs (Wilcoxon test). Ohnolog promoters were compared by computing the Spearman correlation of the TFBS counts.*

### Ohnologs with liver-specific adaptive gains in expression

***Supplementary table 5. The ohnolog pairs where one copy had evolved liver-specific adaptive gains in expression following WGD.***

| Gene up ID | EVE p-value | Tau up | Gene cons ID | Tau cons | Symbol | Description |
|---|---|---|---|---|---|---|
| 106599366 | 1.30E-03 | 0.62 | 106569256 | 0.35 | | zinc finger SWIM domain-containing protein 5-like |
| 100380336 | 9.99E-03 | 0.74 | 106606370 | 0.65 | coda1 | Collagen alpha-1XIII chain |
| 106586620 | 1.32E-03 | 0.65 | 106582100 | 0.6 | | anosmin-1-like |
| 106575074 | 5.00E-03 | 0.68 | 106574305 | 0.81 | | leucine-rich repeat-containing protein 58-like |
| 106611046 | 2.96E-02 | 0.86 | 106591116 | 0.87 | | feline leukemia virus subgroup C receptor-related protein 2-like |
| 106611770 | 1.30E-05 | 0.88 | 106604699 | 0.79 | | GDNF family receptor alpha-4-like |
| 106588972 | 8.67E-05 | 0.66 | 106570416 | 0.49 | | WNT1-inducible-signaling pathway protein 1-like |
| 106565672 | 3.66E-02 | 0.75 | 106583072 | 0.81 | | epidermal growth factor receptor kinase substrate 8-like protein 3 |
| 106583442 | 1.46E-02 | 0.72 | 106565162 | 0.8 | | uncharacterized |
| 106607531 | 1.72E-02 | 0.87 | 106600959 | 0.98 | | uncharacterized |
| 106607884 | 1.16E-02 | 0.62 | 106571468 | 0.66 | disp1 | protein dispatched homolog 1-like |
| 106603096 | 3.63E-02 | 0.66 | 106563420 | 0.56 | chrdl2 | chordin-like 2 |
| 106565918 | 3.32E-02 | 0.77 | 100380427 | 0.58 | errfi | ERBB receptor feedback inhibitor 1-like |
| 106585185 | 3.06E-04 | 0.8 | 106580615 | 0.44 | slc35d2 | UDP-N-acetylglucosamine/UDP-glucose/GDP-mannose transporter-like |
| 100196581 | 1.74E-02 | 0.69 | 106571544 | 0.72 | batf3 | basic leucine zipper transcription factor, ATF-like 3 |
| 106568960 | 2.56E-02 | 0.77 | 106598962 | 0.79 | ocel1 | MARVEL domain-containing protein 2-like |
| 106613481 | 4.53E-03 | 0.71 | 106574163 | 0.75 | | nuclear receptor subfamily 5 group A member 2 |
| 106582073 | 2.33E-02 | 0.74 | 100195006 | 0.72 | pla1a | phospholipase A1 member A-like |

| | | | | | | |
|---|---|---|---|---|---|---|
| 106563436 | 1.49E-03 | 0.68 | 106603109 | 0.44 | | interleukin-13 receptor subunit alpha-2-like |
| 100192340 | 4.10E-02 | 0.86 | 100136433 | 0.43 | elvol5a | polyunsaturated fatty acid elongase elovl5b |
| 106611625 | 1.57E-03 | 0.62 | 106604591 | 0.46 | fgf2 | fibroblast growth factor 2 (basic) |
| 106584157 | 1.66E-02 | 0.75 | 106560308 | 0.47 | | zinc finger protein 644-like |
| 106562145 | 2.15E-02 | 0.86 | 106587237 | 0.83 | | immunoglobulin superfamily DCC subclass member 4-like |
| 106585186 | 8.68E-03 | 0.61 | 106580614 | 0.57 | znf367 | zinc finger protein 367-like |
| 106607938 | 2.42E-02 | 0.67 | 106571501 | 0.45 | | nuclear receptor coactivator 7-like |
| 106579910 | 4.03E-02 | 0.89 | 106585530 | 0.9 | | transmembrane protein 132D-like |
| 106603067 | 1.18E-02 | 0.82 | 106563583 | 0.92 | | MAP7 domain-containing protein 2-like |
| 106611011 | 1.60E-02 | 0.72 | 106595839 | 0.75 | | uncharacterized |
| 100196878 | 1.76E-03 | 0.67 | 106609893 | 0.51 | m4a12 | membrane-spanning 4-domains subfamily A member 12 |
| 106584545 | 2.30E-02 | 0.92 | 106605216 | 0.66 | | uncharacterized |

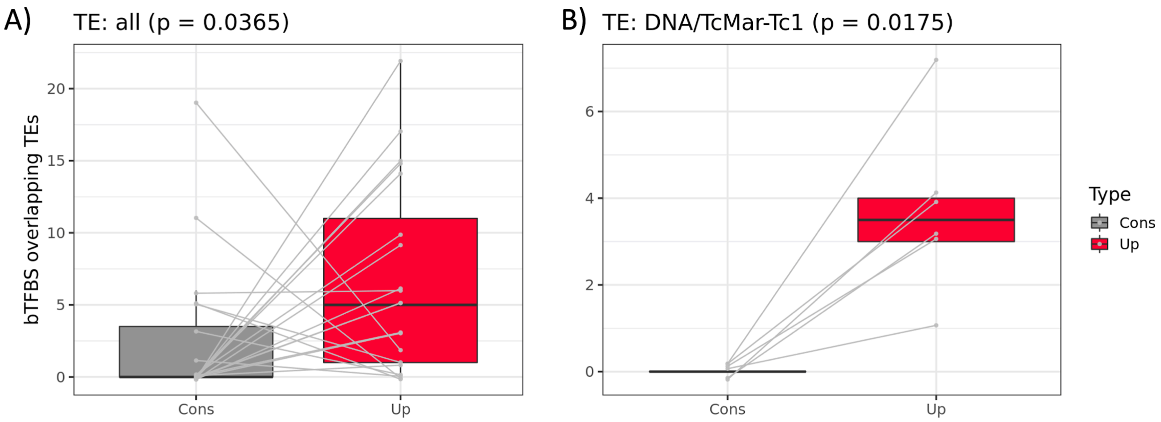


***Supplementary figure 17: TE-bTFBSs overlap**. Number of bTFBSs that overlap TEs in copies with a liver specific gain in expression (Up) and their conserved partner (Cons). Grey lines link ohnolog-pairs, with jitter added to separate identical data points. P-values were obtained by one-sided paired Wilcoxon tests.*

***Supplementary table 6. bTFBSs overlapping the TE superfamily TIR TC1-Mariner.***

| ***TF_Motif*** | ***Gene*** | ***Descr*** |
|---|---|---|
| *HNF4A_MA0114.4* | *106599366* | *zinc finger SWIM domain-containing protein 5-like* |
| *HNF4G_MA0484.2* | *106599366* | *zinc finger SWIM domain-containing protein 5-like* |
| *KLF10_MA1511.1* | *106599366* | *zinc finger SWIM domain-containing protein 5-like* |
| *KLF15_MA1513.1* | *106599366* | *zinc finger SWIM domain-containing protein 5-like* |
| *TBX1_MA0805.1* | *106607938* | *nuclear receptor coactivator 7-like* |
| *PBX3_MA1114.1* | *106607938* | *nuclear receptor coactivator 7-like* |
| *Nr1h3Rxra_MA0494.1* | *106607938* | *nuclear receptor coactivator 7-like* |
| *MAFF_MA0495.3* | *106611011* | *uncharacterized* |
| *GATA6_MA1104.2* | *106613481* | *nuclear receptor subfamily 5 group A member 2* |
| *Arid3b_MA0601.1* | *106613481* | *nuclear receptor subfamily 5 group A member 2* |
| *KLF10_MA1511.1* | *106613481* | *nuclear receptor subfamily 5 group A member 2* |
| *RXRGvar.2_MA1556.1* | *106613481* | *nuclear receptor subfamily 5 group A member 2* |
| *RXRGvar.2_MA1556.1* | *106613481* | *nuclear receptor subfamily 5 group A member 2* |
| *PLAG1_MA0163.1* | *106613481* | *nuclear receptor subfamily 5 group A member 2* |
| *NR1H4RXRA_MA1146.1* | *106613481* | *nuclear receptor subfamily 5 group A member 2* |
| *ELF3_MA0640.2* | *106575074* | *leucine-rich repeat-containing protein 58-like* |
| *EHF_MA0598.3* | *106575074* | *leucine-rich repeat-containing protein 58-like* |
| *TFCP2_MA0145.3* | *106575074* | *leucine-rich repeat-containing protein 58-like* |
| *MAFK_MA0496.3* | *106586620* | *anosmin-1-like* |
| *HNF1A_MA0046.2* | *106586620* | *anosmin-1-like* |
| *HNF1B_MA0153.2* | *106586620* | *anosmin-1-like* |
| *Arid3a_MA0151.1* | *106586620* | *anosmin-1-like* |
